# Alternate recognition by dengue protease: Proteolytic and binding assays provide functional evidence beyond an induced-fit

**DOI:** 10.1101/2024.04.15.589505

**Authors:** Mira A. M. Behnam, Christian D. Klein

## Abstract

Proteases are key enzymes in viral replication, and interfering with these targets is the basis for therapeutic interventions. We previously introduced a hypothesis about conformational selection in the protease of dengue virus and related flaviviruses, based on conformational plasticity noted in X-ray structures. The present work presents the first functional evidence for alternate recognition by the dengue protease, in a mechanism based primarily on conformational selection rather than induced-fit. Recognition of distinct substrates and inhibitors in proteolytic and binding assays varies to a different extent, depending on factors known to influence the dengue protease structure such as pH and salinity. Furthermore, the buffer type and temperature cause a change in binding, proteolysis, or inhibition behavior. Using representative inhibitors with distinct structural scaffolds, we identify two contrasting binding profiles to dengue protease. Noticeable effects are observed in the binding assay upon inclusion of a non-ionic detergent in comparison to the proteolytic assay. The findings highlight the impact of the selection of testing conditions on the observed ligand affinity or inhibitory potency. From a broader scope, the dengue protease presents an example, where the induced-fit paradigm appears insufficient to explain binding events with the biological target. Furthermore, this protein reveals the complexity of comparing or combining biochemical assay data obtained under different conditions. This can be particularly critical for artificial intelligence (AI) approaches in drug discovery that rely on large datasets compiled from different sources.

**HIGHLIGHTS:**

- Buffer type, pH, salt, and temperature influence ligand recognition.
- Experimental conditions in binding and proteolytic assays affect the obtained data.
- Ligand recognition in DENV protease involves mainly conformational selection.

**Graphic for table of contents:** 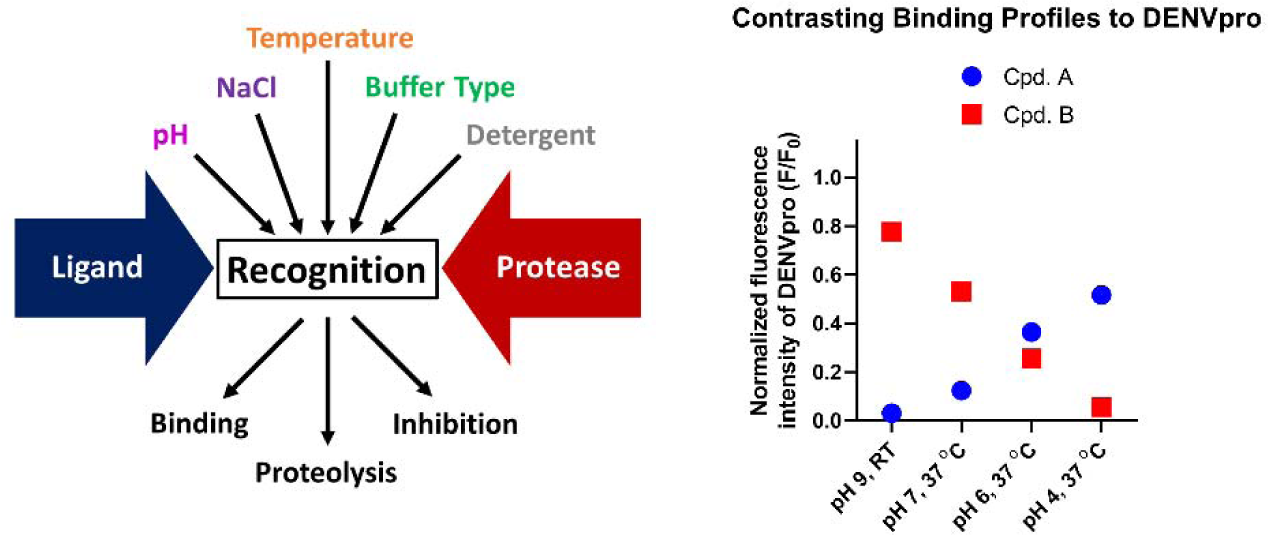

## 1. INTRODUCTION

Proteases from coronaviruses and flaviviruses are important enzymes due to their vital role in viral replication and pathogenesis. These complex targets are responsible for the spatially and temporally controlled processing of the viral polyprotein, which takes place in specific cellular locations and in a distinct order of cleavage events.^1^ On the clinical level, inhibition of viral proteases is a successful treatment approach for viral infections such as HIV. A SARS-CoV-2 main protease (M^pro^) inhibitor is currently approved.^2^ Despite the identification of promising candidates, no protease inhibitor was approved for flaviviruses to date. The development of effective antiviral strategies against viral proteases depends strongly on our understanding of how these proteins function. A key aspect to consider is their molecular recognition of substrates and inhibitors, as it provides the basis for the design of assay systems, enabling the identification of promising hits.

Dengue virus protease (DENVpro), as other proteases from flaviviruses, such as West Nile, Zika, Japanese Encephalitis, and yellow fever viruses, is a complex of the non-structural (NS) proteins NS3 and the hydrophilic core region of NS2B. It recognizes natural cleavage sites possessing one or more basic residues. Structurally, the interaction of the NS2B C-terminus (NS2B-C) with NS3 alters the non-prime sites and the newly identified subpocket B (Figure 1A),^3^ and was initially considered by earlier work as the sole criterion for a functionally active enzyme based on an induced-fit model.^4^

**Figure 1.**
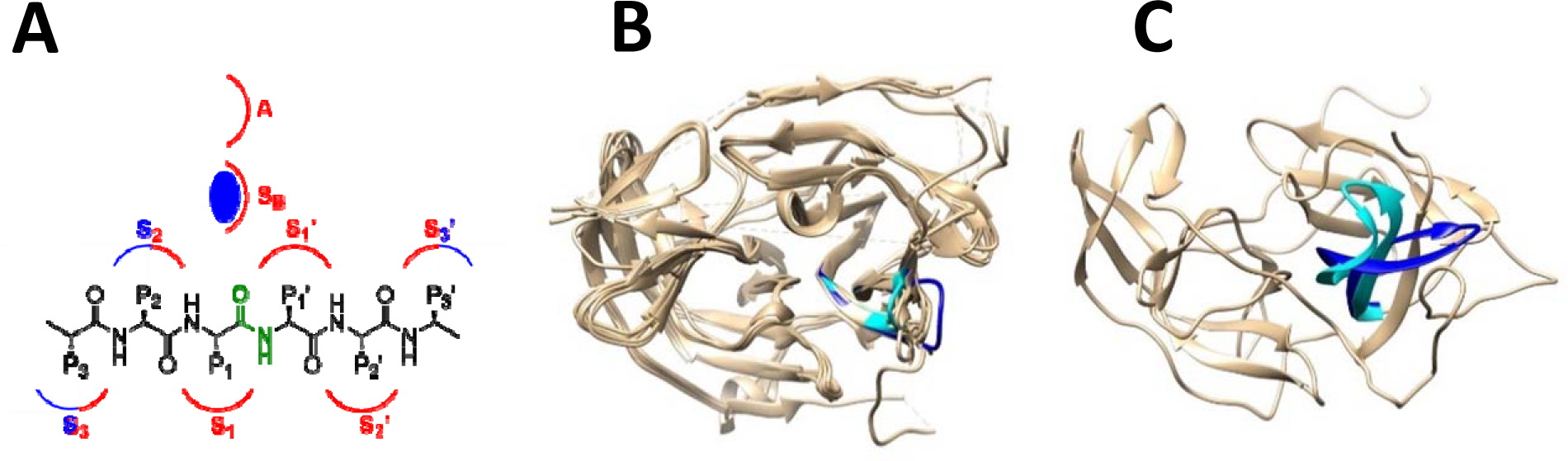
Binding site of the flavivirus protease. (**A**) Schematic representation of the variation in the active site of the flavivirus protease caused by the binding of the NS2B to NS3. Subpockets are indicated with the letter S, S_B_ is subpocket B, A is the allosteric site. S_1_, S_2_, and S_3_ represent the non-prime subpockets, while S_1_’, S_2_’, and S_3_’ represent the prime subpockets. P letters refer to the side chains of amino acids of the substrate. Cleavable amide bond is colored in green. NS2B is shown in blue, and NS3 in red. The blue sphere shows the binding of NS2B-C to S_B_. (**B**) Overlay of X-ray structures of the proteases from WNV (PDB code: 5IDK), ZIKV (PDB code: 5LC0), DENV2 (PDB code: 2FOM), and DENV3 (PDB code: 3U1I). In the overlay, plasticity of the protease is observed beyond the non-prime subpockets, as in the loop extending from residue 21 to 40, otherwise referred to as 30s loop. (**C**) Overlay of NMR structures of DENV2 protease (PDB codes: 2M9P and 2M9Q), shows higher flexibility in 30s loop and a distinct orientation of NS2B C-terminus. Indicated in cyan and blue are the extent of dynamics observed in the 30s loop or residues 21-40 of NS3. Protein chains are colored in tan, in B: 30s loop in 5LC0 chain C in cyan, and in 5LC0 chain A in blue, in C: 30s loop in 2M9Q in cyan and in 2M9P in blue. Overlay is generated by UCSF Chimera.^33^

The induced-fit concept generally governs the mechanism of action of proteases, where ligand binding is responsible for obtaining an active enzyme through conformational changes. Through an in-depth structural analysis, we noted ligand-independent conformational changes for coronavirus and flavivirus proteases. These are not accounted for in a mechanistic understanding based on the induced-fit model, which can seriously impede structural and functional studies, in addition to drug discovery efforts. We thus introduced a hypothesis based on the conformational selection model with secondary induced-fit events,^3,5^ where ligands are distinctly recognized by different active conformations of the same protein.

Basché, Schirmeister and colleagues could distinguish between the induced-fit and conformational selection models for ligand binding to DENVpro by monitoring the NS2B-C interaction with NS3 in a commonly used buffer (Tris, pH 9 with glycerol and CHAPS as additives).^6^ The study showed that the ligand shifts the equilibrium of pre-existing conformations in solution, as proposed by a conformational selection model.^7,8^

The protein conformational landscape in solution is influenced by extrinsic factors, such as pH, temperature, ionic strength, and certain chemical species.^9,10^ As a result, an important implication of the conformational selection model is the dependency of apparent ligand binding affinity on these factors due to interaction with alternate conformations.^8–11^

It is important to note that the kinetic distinction between induced-fit and conformational selection in ligand binding to DENVpro or other flaviviral proteases under a given assessment condition may be influenced by the selected experimental environment of the study, resulting in the nature of the binding event being only secondary to the pre-chosen extrinsic factors. Furthermore, variation in the nature of binding events is possible depending on the assessed ligand, the studied protease, and other variables, as recently observed.^12^ This kinetic assessment is better approached by more specialized techniques that are not in the scope of this work.

Based on the induced-fit model, irrespective of the conditions for measurement in structural or functional studies on the isolated protease, the presence of a particular ligand will induce the enzyme conformation required for its recognition and binding. According to this understanding, the choice of testing conditions for the isolated flaviviral protease is not considered of critical relevance in hit selection. The assays on the isolated flaviviral proteases are usually performed at basic pH, resulting in a high proteolytic efficacy towards tribasic or dibasic substrates. The exact composition of the buffer used and experimental set-up for assessing the catalytic activity varies according to the preference of different research groups in the field. Thus far, mainly the detergent type received attention as a factor that influences the inhibitory activity of selected compounds against the flaviviral protease and other enzymes.^13–15^ Non-ionic detergents, as opposed to the zwitterionic detergent CHAPS, masked the inhibition of certain scaffolds,^14,16^ a point, which we could also confirm in our work.^15^ Triton X-100 was shown to interfere with specific binding of inhibitors to DENVpro^17^ at concentrations where limited solubility and aggregation could be excluded by increasing the DMSO concentration and dynamic light scattering.^16^

The proteolytic assay under standard conditions (Tris buffer, pH 9) with recombinant protease does, for some compound classes, not have a reliable predictive value for antiviral activity. Poor correlation of biochemical data to results of cellular assays that measure viral replication^15,17–19^ was found previously. In contrast, a recently developed luciferase-based DENV2 protease reporter assay in HeLa cells (DENV2proHeLa)^20^ provides high correlation for compound inhibitory activity with the titer reduction assay, which constitutes the gold standard in antiviral drug discovery. The origin of the discrepancy of the results in the reporter vs. biochemical assays could not be traced back to pharmacokinetic factors alone, prompting thus an investigation of molecular recognition by DENV2pro.

We obtained the first results revealing the limitation of the induced-fit paradigm for ligand recognition by SARS-CoV-2 M^pro^. Interestingly, we identified sensitivity of inhibition by boceprevir to testing conditions, with a remarkable 4-fold variation of potency by altering buffer composition.^21^ Furthermore, the reported peak pH for processing by SARS-CoV-2 M^pro^ appears dependent on the used substrate and buffer components, as in our hands, different pH profiles are obtained.^21,22^ The tendency can be interpreted by conformational plasticity of M^pro^.^5^

Similar structural heterogeneity was noted for the flavivirus protease.^3,23^ X-ray crystallography studies for the binding mode of a highly affine peptide-boronic acid, developed in our group, revealed conformational changes outside the non-prime subpockets, independent of the NS2B-C dynamics (PDB: 5LC0 and 5IDK).^23,24^ Specific ligand–protein interaction in solution was also validated by NMR.^25^ Furthermore, examples of alternate conformations with potential for catalytic competence are seen in other structures elucidated by X-ray crystallography (for example, PDB: 5GXJ)^26^ or NMR (PDB: 2M9P and 2M9Q).^27^ However, to date, the influence on the functional recognition by the flaviviral protease remains unexplored.

While crystallographic data hint towards plasticity of the flaviviral protease, also in the absence of a ligand, an influence of crystal packing and contacts, besides other non-physiological factors cannot be excluded for this method. NMR studies also report conformational changes in the ligand-unbound protease depending on the pH and salt concentration.^28^ Furthermore, observations were reported for differences in NMR signal dispersion at 37 °C vs. 25 °C,^29^ and in MES vs. Tris buffer at the same pH.^30^ However, in some cases, the low resolution of X-ray structures or limited sensitivity of NMR hinders the identification of the catalytic competence of a particular conformation through the orientation of the oxyanion hole. Furthermore, crystal structures often do not represent the actual conformational distribution in solution. A case for DENVpro was observed, where NMR depicts different conformations from those obtained by X-ray crystallography (Figure 1B and 1C).^27^ Thus, it is not clear how the processing of a substrate or the binding of a ligand changes under conditions for structural studies, which may elicit conformational changes. It is possible as well that structural techniques cannot capture the entire conformational landscape of a protein to explain potential changes in ligand recognition in biochemical assays. Functional and binding assays are performed at low protein concentrations, as opposed to commonly used protocols for X-ray crystallization and NMR studies, which require a concentration range of 150 *µ*M or higher.^31^ Concentration-dependent changes in protein conformation favor compact structures, promote association of subunits and the formation of higher-order structures with the increase in concentration.^32^

We therefore decided to explore whether the observed structural plasticity can have a functional significance in proteolytic or binding assays that use the isolated protein, by studying the influence of testing conditions on the cleavage and recognition of different ligands, substrates and inhibitors, by DENVpro. Our focus is directed at factors, previously reported to influence the protease structure, especially from studies in solution. We here use the terminology “proteolytic assay” to indicate an experiment that studies the catalytic activity, and “binding assay” to denote a non-catalytic ligand binding assay which was recently described by us^34^ (Behnam, Basché, Klein; Analytical Biochemistry 2023; manuscript accepted for publication; unformatted draft is provided as review-only supplementary material).

In the present study, we demonstrate that DENVpro, similar to SARS-CoV-2 M^pro^, possesses variable molecular recognition profiles for substrates and inhibitors depending on experimental factors of known influence on the protease conformation. Furthermore, an unexpected temperature-sensitivity is observed for DENVpro in ligand binding and proteolytic processing, the latter being both ligand- and buffer-specific. The sum of our findings presents functional evidence into a mechanism of action of DENVpro that cannot be explained solely by induced-fit, and – in continuation of previous work by others and us – that assay results depend on the selection of assay conditions. The results support the conformational selection paradigm, and allow a conceptual advancement in the understanding of viral proteases with broad implications on the study and targeting of this enzyme group. With respect to the growing trends in using artificial intelligence models in drug discovery, the identified factors behind the variability of assay results in this case study call for caution when extended datasets are assembled from the literature.

## 2. MATERIAL AND METHODS

### 2.1. Binding mode and structural analysis

Model for the binding mode of compound **1** and depiction of conformational dynamics in the flavivirus protease were performed with UCSF Chimera version 1.15rc.^33^ PDB files with more than one unit of the protein were split to individual units that are saved separately. For the overlay, a structural alignment was performed with the “Match Maker” tool in Chimera. Default settings were used for visualization. For modeling the binding of compound **1** in the active site, the structure of the co-crystallized inhibitor **CN-716** with WNV and ZIKV proteases was used in an overlay with DENV3 protease structure. The side chain for the P2 residue in **CN-716** was exchanged for that of compound **1** using the build model tool and the boronate ester with glycerol was removed.

### 2.2. Expression and purification of DENV2 protease

DENV (serotype 2) NS2B−NS3 protease construct possessing a glycine serine GGGGSGGGG linker, which covalently connects the NS3 protease and NS2B hydrophilic core domain was used for this study, and was prepared by C. Steuer. Expression and purification of the His_6_-tagged protein was performed as previously described^35^ with some modifications. After cell disruption and centrifugation, DNAseI (Sigma-Aldrich) was added to the supernatant and purification was carried by Ni^2+^-affinity chromatography using NTA-agarose column (Chelating Sepharose TM, Fast Flow, GE Healthcare). The sample was loaded and washed with 500 mL lysis buffer (25 mM Tris pH 7.9, 0.5 M NaCl, 5 mM imidazole, 5% glycerol), then the target protein was eluted with the elution buffer (25 mM Tris pH 7.9, 0.5 M NaCl, 250 mM imidazole, 5% glycerol), fractions were collected and checked by OD measurement and SDS-page. The identity of the eluted target protein was confirmed by MALDIToF-MS and HR-ESI using LC/MS. (LC/MS method is described below) For the removal of the His_6_-tag, Thrombin CleanCleave^TM^ Kit containing immobilized thrombin was used. Purification of the non-tagged protein was performed by a second passage through Ni^2+^-NTA agarose column, followed by size exclusion chromatography using an HiLoad^TM^ 16/60 Superdex^TM^ 200 pg column and a running buffer (50 mM Tris, 150 mM NaCl, pH 8). Fractions with pure protein were concentrated, buffer was exchanged to PBS and mixed with 50% glycerol. Aliquots were flash frozen in liquid nitrogen and stored at −80 °C until further use.

### 2.3. Proteolytic assays

DENV2 protease functional characterization and inhibition assays were performed on a BMG Labtech FLUOstar OMEGA microtiter plate reader. Buffers were prepared with a concentration of 50 mM. Buffers generally included no additives, unless otherwise mentioned. For buffer containing CHAPS, its concentration is 1 mM. Compounds and substrates were prepared as 10 mM stocks in DMSO or 100 mM stocks in DMSO for substrate **18** and *p*NA. Predilutions were performed in transparent 96-well U-bottom plates (Greiner Bio-One, Germany). Final assay volume is 100 _μ_l per well. All experiments were performed in duplicates and averaged. Results were analyzed using GraphPad Prism 8.0.

Fluorescence proteolytic assays were measured as continuous assays in black 96-well V-bottom plates (Greiner Bio-One, Germany). For FRET substrates, excitation and emission wavelengths of 330 nm and 430 nm, respectively were used. For AMC-based substrates, excitation and emission wavelengths of 355 nm and 460 nm, respectively were used. Enzyme final concentration is 100 nM, unless otherwise specified, substrate final concentration is 50 _μ_M. Control experiments in the absence of the enzyme were measured for representative substrates in selected buffers, and in the pH range 7–10 confirming stability. Two protocols were used for assessing the influence of temperature on substrate processing: in the first protocol, the enzyme and all assay solutions are kept on ice, and the assay is measured at RT; in the second protocol, the enzyme and all assay solutions are kept at 37 °C, and the assay is measured at 37 °C. The enzymatic activity was determined as slope per second (relative fluorescence units per second, RFU/s) and monitored for 15 min. IC_50_ determination were performed as previously described^20^ with modification of the used buffer. The incubation of the compounds with the protease was either conducted at RT or at 37 °C. Eight concentrations of the compounds were used covering the range 0–100 *µ*M or 0–50 *µ*M with a final concentration of 1% DMSO to avoid potential precipitation of the compounds. Highly active compounds were assayed in the range 0–5 *µ*M. Percentage inhibition was calculated relative to a positive control (without the inhibitor). Kinetic characterization of representive FRET or AMC-based substrates was carried out comparably to previous protocols.^36,37^ Eight concentrations of the substrates were used in the range of 0–200 *µ*M. The slope for the enzymatic reaction in relative fluorescence units per second (RFU/s) was corrected for inner-filter effect, and using a calibration curve for the fluorophore emission signal to concentration (0-10 *µ*M concentration), the reaction rate was obtained.

Chromogenic proteolytic assays with *p*NA-based substrates were performed as end-point measurements in transparent 96-well flat-bottom plates (Greiner Bio-One, Germany). For substrate **18**, 5 *µ*M enzyme concentration was used and 2.5 mM substrate concentration due to the weak reaction. For substrate **19**, 200 nM enzyme concentration was used and 1 mM substrate concentration. After addition of the substrate, the sample was incubated for 24 hr then measured at 405 nm as absorbance wavelength, in addition to a reference wavelength at 700 nm to monitor any turbidity. Longer incubation for 48 hr required the addition of glycerol (5-20%) and was not further assessed. To prevent solution evaporation, the plate was covered with AeraSeal film (Sigma Aldrich), which was removed before measurement. Control experiments without the enzyme were measured under the same conditions and were used to correct the results.

Fluorescence or absorbance of references such as 2-Abz, AMC, or *p*NA were performed in the same buffers but in the absence of the enzyme.

### 2.4. Binding assays

DENV2 protease binding and competition assays were performed on a BMG Labtech FLUOstar OMEGA microtiter plate reader, in black 96-well V-bottom plates (Greiner Bio-One, Germany). Excitation and emission wavelengths were 295 nm and 350 nm, respectively, as previously described.^34^ Buffers were prepared with 50 mM concentration, and contained 1 mM CHAPS. Compounds were prepared as 10 mM stocks in DMSO or 1 mM stock in water for aprotinin. Predilutions were performed in transparent 96-well U-bottom plates (Greiner Bio-One, Germany). Final concentration in the assay is 200 nM for DENV2pro or 1 *µ*M for NATA, and final assay volume is 100 _μ_l per well. As additional precaution against precipitation of compounds, a final concentration of 1% or 2% DMSO was used in the assay. The following measurement conditions were assessed: Tris-CHAPS buffer, pH 9, RT, His_6_-DENV2pro; phosphate-CHAPS, pH 7, 0 °C, phosphate-CHAPS, pH 7, 37 °C, non-tagged DENV2pro; phosphate-CHAPS, pH 6, 37 °C, non-tagged DENV2pro; acetate-CHAPS, pH 4, 37 °C, non-tagged DENV2pro; and citrate-CHAPS, pH 4, 37 °C, non-tagged DENV2pro. Samples were incubated 1 hr before measurement, background signals were subtracted from recorded fluorescence intensity and the data was corrected using correction factors derived from NATA experiments. Finally, the obtained fluorescence intensity was normalized relative to that of the protease. Correction factors were obtained by dividing the fluorescence intensity of 1 *µ*M NATA in the absence of the ligand, by that in the presence of the ligand. For competition experiments, the displacement was corrected for the autofluorescence of the competing ligand, by measuring the fluorescence of the protease with the ligand in the absence of the quencher. To exclude a difference in solution turbidity under different measurement conditions, absorbance at 700 nm was measured on a BMG Labtech FLUOstar OMEGA microtiter plate reader. For this purpose, after incubation for 1 hr and measurement of the binding assay, 80 *µ*l of the assayed solution were transferred to a transparent 96-well flat-bottom plate (Greiner Bio-One, Germany). *K*_d_ and IC_50_ determinations for compounds **20**, **21**, and **23** were carried out using 1 hr incubation. Eight concentrations were used in the range of 0–100 *µ*M for **20** and **21**, or 0–25 *µ*M for **23**. For the binding curves, background signals were subtracted from all data, recorded fluorescence quenching was corrected for inner filter effects based on experiments with NATA instead of the protease enzyme, and the data was fitted as previously described.^38^ All experiments were performed in duplicates and averaged. Results were analyzed using GraphPad Prism 8.0 or OriginPro 2021.

### 2.5. LC/MS degradation studies

Degradation of compound **2** by DENV2 protease was assessed in a comparable manner to previously described procedures.^39^ Compound **2** in a concentration of 100 _μ_M was incubated with DENV2 protease (final concentration 3 _μ_M) in the indicated assay buffer and conditions for 24 hr. A volume of 20 _μ_l of incubated solution was subjected to LC/MS analysis. The measurements were performed on an Agilent 1200 series HPLC device in combination with an ESI-MS micrOTOF-QII instrument (Bruker Daltonik, Germany) using Reprosil-Pur ODS-3, Dr. Maisch GmbH, Germany, 3 _μ_m, 50 × 2 mm HPLC column. The method was as follows: eluent A, water (0.1% formic acid); eluent B, acetonitrile (0.1% formic acid); 0−4.0 min, gradient 10% B to 95% B; 4.1−8.0 min, isocratic 95% B; 8.1−13.0 min, isocratic 10% B; flow rate, 0.3 ml/min. HRMS measurements were performed in positive mode.

The protease mass could be monitored in each run by deconvolution of the MS results. The intact mass of the protein did not show relevant variation between different assay buffers and conditions.

### 2.6. Reagents and solvents

Chemicals used for synthesis were obtained from Sigma-Aldrich (Germany), Alfa Aesar (Germany), TCI Europe (Belgium), and were of analytical grade. All solvents were used as obtained from the commercial sources. The solid-phase resins, protected amino acids, coupling reagents were purchased from Bachem (Switzerland), Iris Biotech (Germany), Orpegen (Germany), and Carbolution Chemicals (Germany).

### 2.7. NMR analysis

NMR spectra for compounds **1**, **20** and **21** were recorded in D_2_O or DMSO-*d*_6_ NMR on Varian NMR instruments at 300 or 500 MHz, 300 K. Chemical shifts (δ) are given in parts per million (ppm) in reference to residuals of nondeuterated solvent as internal standard for ^1^H NMR: D_2_O: δ = 4.79 ppm and DMSO-*d*_6_: δ = 2.50 ppm. Coupling constants (J) are given in hertz (Hz). Multiplicity is reported as s (singlet), d (doublet), t (triplet) and m (multiplet), respectively.

### 2.8. HPLC analysis

HR-ESI mass spectra were measured on Bruker microTOF-Q II instrument. Purity of synthesized peptide substrates for DENV2 was determined by HPLC on a Jasco HPLC system with a DAD detector on an RP-18 column (ReproSil-Pur-ODS-3, Dr. Maisch GmbH, Germany, 5 μm, 50 mm × 2 mm) using the following conditions: water (0.1% TFA), eluent B: acetonitrile (0.1% TFA), injection volume: 10 *µ*l, flow rate: 1 ml/min, gradient: 1% B (0.2 min), 100% B (7 min), 100% B (8 min), 1% B (8.1 min), 1% B (10 min). UV-detection was performed at 254 nm. Analysis was carried out using 100 µM samples in water/acetonitrile (1:1). All tested target compounds and substrates were obtained with a purity of at least 95%, unless otherwise indicated.

### 2.9. Synthesis of substrates

Substrate **18** was purchased commercially from Bachem (Switzerland). All other substrates were synthesized using Fmoc-strategy by solid-phase peptide synthesis for FRET substrates, or a combination of solid-phase synthesis with in-solution coupling for AMC- or *p*NA-based substrates. Substrate **6** is previously reported for DENV2 protease,^40^ and substrate **13** for ZIKV and WNV proteases.^41^ Substrate **17** is commonly used for DENV2 and ZIKV proteases.

Synthesis of substrates **6-15** was performed on Rink amide resin having an average capacity of 0.65 mmol/g, with minor modifications compared to previously published procedures.^36^ 200 mg Rink amide resin were used for each peptide. 3 equivalents of the protected amino acid or Fmoc-Abz-OH were used for each coupling step. Coupling was carried out using HATU/TMP (3 and 4 equivalents, respectively, relative to Rink amide resin), and Fmoc-cleavage steps were performed with 10% piperidine in DMF. Cleavage solution is TFA:H_2_O:TIPS (95:4:1). Purification was performed by HPLC Äkta purifier system (Cytiva, formerly GE Healthcare) with the following method: RP-18 pre- and main column (Reprospher 100 C-18-DE, Dr. Maisch GmbH, Germany, 5 μm, precolumn 30 × 16 mm, main column 125 × 16 mm); eluent A, water (0.1% TFA); eluent B, methanol (0.1% TFA); 0−2.5 min, 10% B; 2.6−23.5 min, gradient 10% B to 100% B; 23.6−26.0 min, 100% B; 26.1−30.0 min, 10% B; flow rate, 8 mL/min; λ = 214, 254, and 280 nm. After lyophilization, peptides were obtained as yellow solid.

For the substrates **16-17** and **19**, 2-chlorotrityl chloride resin (CTC) had an average capacity of 1 mmol/g was used to synthesis the first part of the structure with the sequence Bz-nK(Boc)R(Pbf)-OH or (*t*-Bu) acetic acid-GR(Pbf)-OH. Synthesis took place with minor modifications compared to previously published procedure.^37^ 500 mg CTC resin were used for each sequence. Loading of the first amino acid (3 equivalents) was performed overnight using TMP (3 equivalents relative to the resin). Resin capping solution is DCM:MeOH:TMP (80:15:5). Coupling was carried out using HATU/TMP (3 and 4 equivalents, respectively, relative to Rink amide resin), and Fmoc-cleavage steps were performed with 10% piperidine in DMF. Cleavage solution is 1% TFA in DCM.

The crude product was obtained as white solid and used for the following step of in-solution coupling to Arg-AMC or Arg-*p*NA, as described below.

Substrate **16**: (*t*-Bu) acetic acid-GR(Pbf)-OH, intermediate **Ia**, was obtained as white solid (49 mg, 16.9% yield). In a small flask, 41 mg (0.07 mmol, 1 eqv.) of intermediate **Ia**, 31 mg (0.077 mmol, MW=404.292, 1.1 eqv.) Arg-AMC.2HCl, 29 mg (0.077 mmol, MW = 380.23, 1.1 eqv.) HATU, 93 ul (0.7 mmol, MW = 121.8, 10 eqv.) were added with 2 ml DMF. Reaction was stirred at room temperature for 48 hr. Solution was acidify with 1N HCl, diluted with water, and solvents were removed under vacuum at 60 degrees. Purification was performed by HPLC Äkta purifier system as described above. After lyophilization, substrate **16** was obtained as white solid.

Substrate **17**: Bz-nK(Boc)R(Pbf)-OH, intermediate **Ib**, was obtained as white solid (83 mg, 18.8% yield). In a small flask, 50 mg (0.057 mmol, 1 eqv.) of intermediate **Ib**, 25 mg (0.063 mmol, MW=404.292, 1.1 eqv.) Arg-AMC.2HCl, 24 mg (0.077 mmol, MW = 380.23, 1.1 eqv.) HATU, 28 ul (0.21 mmol, MW = 121.8, 3.7 eqv.) were added with 2 ml DMF. Reaction was stirred at room temperature for 24 hr. Solution was acidify with 1N HCl, diluted with water, and solvents were removed under vacuum at 60 degrees. Purification was performed by HPLC Äkta purifier system as described above. After lyophilization, substrate **17** was obtained as white solid. NMR for the final product: ^1^H NMR (500 MHz, CD_3_OD) δ 8.41 (d, *J* = 7.4 Hz, 0H), 8.36 (d, *J* = 7.4 Hz, 1H), 8.22 (d, *J* = 6.7 Hz, 1H), 7.89 (d, *J* = 2.0 Hz, 1H), 7.87 (d, *J* = 7.2 Hz, 2H), 7.72 (d, *J* = 8.7 Hz, 1H), 7.56 (t, *J* = 7.4 Hz, 1H), 7.53 – 7.44 (m, 3H), 6.26 (s, 1H), 4.55 – 4.46 (m, 1H), 4.45 – 4.32 (m, 3H), 3.24 (t, *J* = 7.1 Hz, 2H), 3.19 (t, *J* = 7.0 Hz, 2H), 2.94 (t, *J* = 7.4 Hz, 2H), 2.46 (s, 3H), 2.02 – 1.63 (m, 14H), 1.62 – 1.45 (m, 3H), 1.45 – 1.36 (m, 3H), 0.94 (t, *J* = 7.1 Hz, 3H).

Substrate **19**: (*t*-Bu) acetic acid-GR(Pbf)-OH, intermediate **Ia**, was obtained as white solid (49 mg, 16.9% yield). In a small flask, 41 mg (0.07 mmol, 1 eqv.) of intermediate **Ia**, 31 mg (0.077 mmol, MW=404.292, 1.1 eqv.) Arg-*p*NA.2HCl, 28 mg (0.077 mmol, MW = 367.24, 1.1 eqv.) HATU, 93 ul (0.7 mmol, MW = 121.8, 10 eqv.) were added with 2 ml DMF. Reaction was stirred at room temperature for 48 hr. Solution was acidify with 1N HCl, diluted with water, and solvents were removed under vacuum at 60 degrees. Purification was performed by HPLC Äkta purifier system as described above. After lyophilization, substrate **19** was obtained as white solid. (refer to the Supporting Information for analytical characterization of synthesized substrates, Table S1)

### 2.10. Synthesis of inhibitors

Synthesis of compound **1** was performed based on previously published procedures^24^ (refer to the Supporting Information for analytical characterization).

Compounds **2, 20-23** were synthesized as previously described (refer to the Supporting Information for analytical characterization of synthesized inhibitors) with a modified procedure, where HATU/TMP were used for coupling instead of HATU/DIPEA.^42,43^ Compounds **3-5** were provided by Nikos Kühl and Christian Gege.^15,20,35^ Compounds **24** and **25** were purchased. ARDP0006 was purchased from Sigma-Aldrich and nitazoxanide was purchased from TCI. Compound **26** was provided by Yuxing Deng based on previously reported^17^ synthetic procedures.

## 3. RESULTS AND DISCUSSION

### 3.1. Recognition in the proteolytic assay

Representative inhibitors from different structural classes that show potent inhibition in cellular assays were selected (Figure 2A).^15,20,35,42^ Validation of included compounds was previously performed in titer reduction assays, in addition to a recently developed luciferase-based DENV2 protease reporter assay in HeLa cells (DENV2proHeLa).^20^ The expected binding mode of **1** in the active site of the protease is depicted in Figure 2B. A set of FRET, AMC, and *p*NA-based substrates were synthesized guided by natural cleavage sequences (Figure 2C). Interestingly, the FRET-based substrate **9** showed comparable kinetic parameters (cf. Supporting Information, Table S1) to the commonly used AMC-based substrate **17** in Tris buffer, pH 9. Based on their high proteolytic processing in Tris buffer, pH 9 (cf. Supporting Information, Figures S1 and S2), both **9** and **17** were further evaluated in different buffer compositions and pH (Figure 2D). This was also performed for the monobasic substrate **18** (Figure 2E) and for the inhibition of compound **2** (Figure 2F). Furthermore, inhibitory activity was evaluated under variable conditions for the reference inhibitors (Table 1).

**Figure 2.**
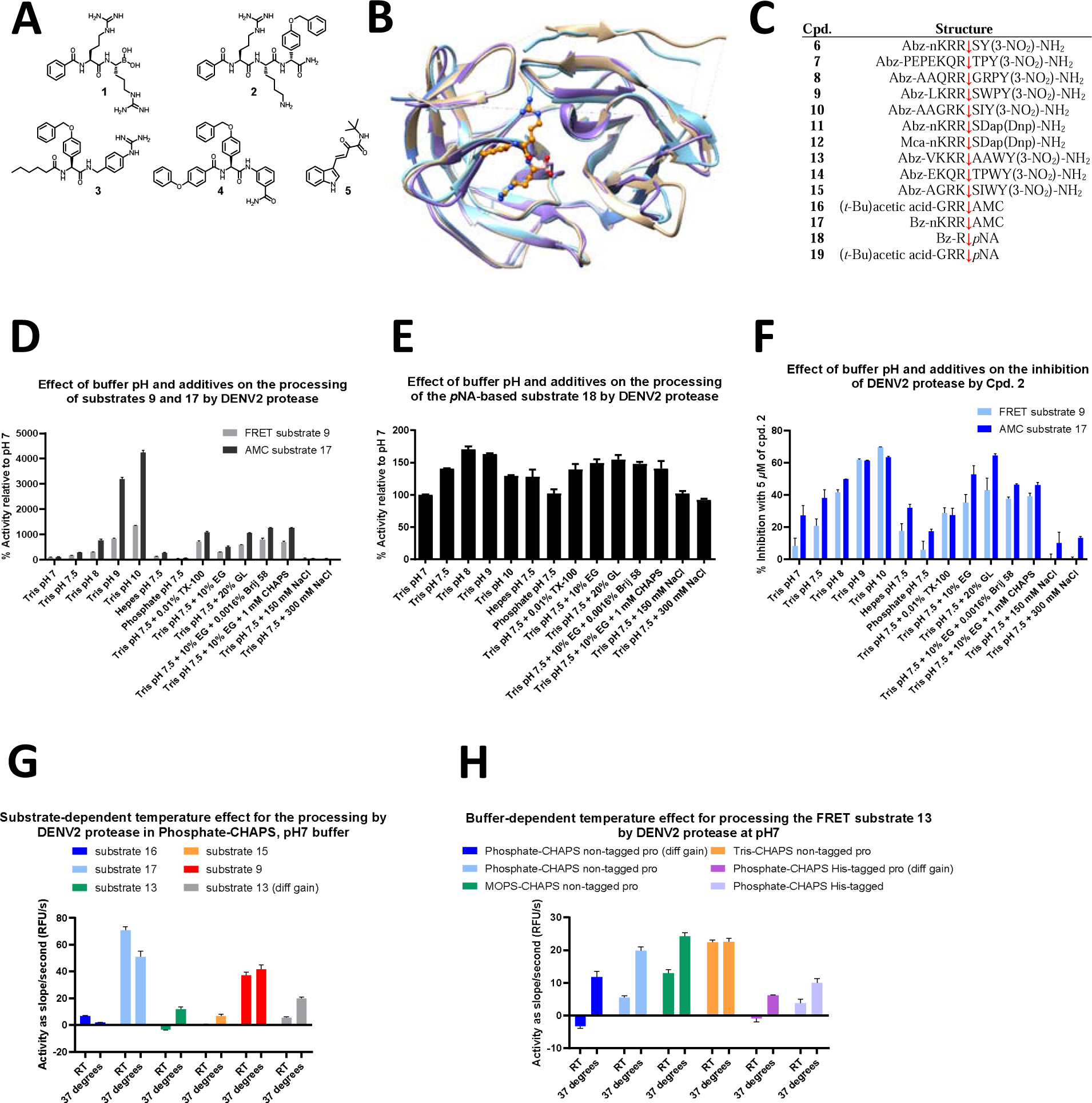
Alternate molecular recognition by DENV2 protease in the proteolytic assay. (**A**) Structure of inhibitors included in this study. (**B**) Model for the binding mode of compound **1** in the active site of the flavivirus protease. Compound **1** is represented in ball and stick format in orange. Overlay of X-ray structures is shown in ribbon format for the NS2B–NS3 protease of DENV3 (PDB: 3U1I chains A and B, light blue), WNV (PDB: 5IDK chain A, tan), and ZIKV (PDB: 5LC0 chain A, light purple) using UCSF Chimera.^33^ (**C**) Substrates used in proteolytic assays. (**D**) Buffer pH and composition influence the processing of the tribasic substrates **9** and **17** by DENV2 protease. EG stands for ethylene glycol and GL for glycerol. (**E**) Buffer pH and composition exert limited effects on the processing of the monobasic substrate **18** by DENV2 protease. (**F**) Inhibition of DENV2 protease by compound **2** (5 *µ*M) shows remarkable sensitivity to buffer pH and additives. (**G**) Temperature sensitivity of the catalytic processing by DENV2 protease is substrate-dependent in phosphate-CHAPS buffer, pH 7. “diff gain” refers to different gain in acquiring the signal. (**H**) Substrate **13** displays buffer and temperature sensitivity for the catalytic processing by DENV2 protease.

**Table 1.** Variation of half-maximal inhibitory concentration (IC_50_) values for compounds and Michaelis-Menten constants (*K*_m_) for substrates under different assay conditions for DENV2 protease. Inhibitory activity as IC_50_ is determined against 50 *µ*M of substrate **17**.

**IC<sub>50</sub> ( μM) under different assay conditions relative to substrate 17**
|  | 50 mM Tris, pH 9, 10% EG, 1 mM CHAPS | 50 mM Tris, pH 7.5, 10% EG, 1 mM CHAPS | 50 mM Phosphate, pH 7.5, 300 mM NaCl, 10% EG, 1 mM CHAPS | 50 mM Phosphate, pH 7.5, 300 mM NaCl, 10% EG, 1 mM CHAPS | 50 mM Phosphate, pH 7, 1 mM CHAPS |
| --- | --- | --- | --- | --- | --- |
|  | His<sub>6</sub>-DENV2 pro RT | His<sub>6</sub>- DENV2 pro RT | His<sub>6</sub>-DENV2 pro RT | Non-tagged DENV2 pro RT | Non-tagged DENV2 pro 37 degrees |
| **1** | 0.68 ± 0.01 | 0.25 ± 0.01 | 0.52 ± 0.02 | 0.52 ± 0.01 | 0.89 ± 0.03 |
| **2** | 1.71 ± 0.17 | 8.37 ± 0.29 | 48.28 ± 2.24 | 38.64 ± 1.45 | 117.8 ± 7.26 |
| **3** | 17.65 ± 2.15 | 22.92 ± 2.95 | 25.62 ± 2.66 | 26.06 ± 1.95 | 12.05 ± 1.15 |
| **4** | 5.37 ± 0.73 | 6.34 ± 0.43 | 7.87 ± 0.81 | 12.57 ± 1.61 | 11.25 ± 1.22 |
| **5** | 37.08 ± 1.29 | 64.36 ± 2.47 | 73.91 ± 8.61 | 71.51 ± 3.64 | 62.31 ± 6.76 |

**K<sub>m</sub> ( μM) of substrate 17 under different assay conditions**
15.86 ± 1.19 90.36 ± 4.57 149.0 ± 13.75 262.4 ± 17.27 75.52 ± 4.93

Of note, the results of in vitro biochemical assays can be influenced not only by protein-related, but also by ligand-related effects. Ligands may also have an effect on the signal readout. In comparison to SARS-CoV-2 M^pro^, the substrates and inhibitors of the flaviviral protease display in general more favorable properties for the purpose of this study. Indications of precipitation were absent under the tested conditions, and control experiments were performed as described (cf. Material and Methods, and Supporting Information) and discussed further.

#### 3.1.1. Proteolytic recognition of DENVpro substrates depends on buffer composition and temperature

Results of functional characterization of different substrates for the DENV2pro reveal distinct proteolytic profiles, which reflect the influence of studied factors on the recognition of particular substrates by DENVpro rather than an effect on the catalytic competence of the enzyme. In fact, the measurement conditions mostly affect just the cleavage of specific substrates. The cleavage of the tribasic FRET or AMC substrates, **9** and **17**, respectively, is highly sensitive for salt concentration and pH (Figure 2D). These factors are known by NMR studies^28^ to weaken the NS2B-C interaction with NS3, which shapes the S2 and S3 subsites. In comparison to Tris pH 7.5, the inclusion of 150 mM or 300 mM NaCl causes a decrease in the proteolytic processing of **9** by 3- or 4-fold, respectively, and of **17** by 5- or 8-fold, respectively. This is not the case for the monobasic Bz-Arg-*p*NA substrate **18**,^44^ which is not expected to occupy subpockets beyond S1 (Figure 2E). The inclusion of 150 mM or 300 mM NaCl lowers the proteolytic processing of **18** by 1.3- or 1.5-fold, respectively, in comparison to Tris pH 7.5. Due to the slow processing of substrate **18**, relatively higher concentrations of the protease and substrate were used than in the case of substrates **9** or **17**. The concentration of the protease remains, however, considerably lower than that commonly used for NMR studies, where concentrations in the range of 0.5 – 0.8 mM are commonly used. The higher protein concentrations in NMR experiments (or in X-ray crystallography) may favor more compact folded states by crowding effects. Control experiments for **18** revealed comparable background signals and non-specific effects on the absorbance between different buffers, and were used to correct the results. Buffer effects on the signal readout by fluorescence or absorbance were absent (cf. Supporting Information, Figures S3, S4, and S5).

Distinct pH profiles are observed for different substrates or buffers, with variation in the steepness from pH 7 to pH 9 or 10 and the peak pH for processing (Figures 2D, 2E, and Supporting Information Figures S6, S7, S8, and S9). For example, the monobasic substrate **18** has a peak pH for processing at pH 8, with a minor decrease in proteolysis at pH 9 (Figure 2E). On the contrary, dibasic (cf. Supporting Information, Figures S6 and S7) or tribasic substrates (Figure 2D and Supporting information Figure S8) are preferentially processed at higher pH values, with a steep increase in proteolytic efficacy from pH 8 to pH 9. An influence of *p*NA on the observed pH profile of **18** could be excluded, as the pH profile of the dibasic *p*NA-based substrate **19** reveals a peak pH of processing at pH 10, similar to the dibasic AMC-based substrate **16**, although with a slight difference in the steepness of the processing from pH 7 to pH 10 (cf. Supporting Information, Figures S6 and S7).

The inclusion of CHAPS in the buffer changes the substrate processing profile relative to the non-ionic detergent Triton X-100 (Supporting Information, Figure S10). Triton X-100 increases substrate cleavage in the absence of CHAPS, but not in its presence. Usually, detergents are added to prevent non-specific effects owing to surface adsorption or aggregation, which may commonly take place in different types of in vitro assays. Observed enhancement in proteolytic processing in the presence of a detergent is often attributed to these general aspects. These molecules may be present in huge excess in comparison to the ligand and were previously co-crystallized in the binding space of other targets. Detergents can as well substantially disrupt protein interactions and conformations. Li *et al*. reported an influence of non-ionic detergents on the interaction between the NS2B N-terminus and NS3.^45^ The interaction between NS3 and NS2B is relatively strong with an equilibrium binding constant below 200 nM in Tris, pH 8.5 (with glycerol and CHAPS as additives).^46^ In the presence of non-ionic detergents, inhibitors identified by the cellular DENV2proHeLa assay did not show activity in the proteolytic assay. If the zwitterionic detergent CHAPS was used, some activity could be observed for these compounds.^15^

In comparison to the pH and NaCl concentration, the buffer type and temperature are not well-studied factors regarding their impact on the flaviviral protease conformation. For DENVpro, temperature and buffer type exert an influence on proteolytic processing, which is substrate-specific. Conducting the assay at 37 °C can increase, decrease, or not affect the slope of the catalytic reaction depending on the assessed substrate and the used buffer (Figures 2G and 2H). The effect is not caused by a buffer-related influence on the readout in the absence of the protease nor by temperature-sensitivity in the signal of fluorophores (cf. Supporting Information, Figures S12 and S13). The variable response to temperature among assessed substrates in the same buffer does not support a general non-specific effect of this factor, for example on solubility or surface adsorption. For substrates **13** and **15**, the temperature sensitivity varies between phosphate buffer, MOPS, or Tris (Figure 2H and Supporting Information, Figure S11). Interestingly, an NMR study reported distinct signals at the same pH (8.5) in MES vs. Tris buffer, which were attributed to different forms of the complex produced by the same ligand with DENVpro.^30^ The most obvious temperature sensitive processing by DENVpro was noted in phosphate buffer for substrate **13**, a previously reported substrate for ZIKVpro.^41^ An influence of the His_6_-tag could be excluded by using the non-tagged DENV2pro. The influence of the increase in temperature on the pH profile of **13** is minimal, similar to substrate **17** (Supporting Information, Figures S8 and S9). A mechanistic model based only on induced-fit does not encompass a dependency of the proteolytic processing by DENVpro on temperature and buffer type, from a protein-based perspective.

Furthermore, the proteolytic processing profile of substrate **13** is remarkable as normally the pH of Tris buffer, in particular, is more sensitive towards temperature changes than that of phosphate buffer. This would indicate that the observed effect on substrate processing is related to the temperature as a factor in itself and is not mediated by a temperature-dependent change in pH. As the basic non-prime amino acids are comparable in the substrates studied here, the effect may be caused by non-basic prime side moieties whose binding tendency is less sensitive towards pH changes.

#### 3.1.2. Assay conditions influence the potency of inhibitors in the proteolytic assay

As previously shown for SARS-CoV-2 M^pro^, different buffers and measurement conditions influence substrate kinetic parameters (cf. Supporting Information, Table S2) and in some cases inhibitor potency (Figure 2F and Table 1). Non-specific or ligand-related effects could be difficult to distinguish from protein recognition in inhibition assays. However, the previously discussed influence of studied factors on the proteolytic processing of substrates supports an explanation from the perspective of the protein. As some inhibitors may be more prone to solubility problems, a higher final concentration of DMSO (1%) was used during the assessment of inhibition. Michaelis-Menten curves did not show deviations that could indicate a tendency of the substrates for precipitation. In addition, the Hill slopes of IC_50_ curves were in support of specific inhibition.

The inhibitory activity of compound **2** displays a comparable influence with variation in buffer composition and pH to the processing of the tribasic substrates **9** and **17** (Figure 2D and 2F). Relative ranking of hits depends on the choice of measurement conditions. For instance, performing the assay in Tris pH 9, RT selects preferentially for compound **2** over **3** and **4**, while in phosphate, 37 °C compounds **3** and **4** outperform **2** (Table 1). The variation in the IC_50_ values of compound **2** serves as an example for the complexity in comparing results of biochemical assay procedures, even when the identity and concentration of the substrate are not changed. This can be demonstrated further through the influence of experimental conditions on the substrate-independent *K*_d_ values (Table 2).

**Table 2.** Dissociation constants and inhibitory activities of compounds **20** and **21** against DENV2 protease. Inhibitory activity as IC_50_ is determined against 50 *µ*M of substrate **17**. IC_50_ values and dissociation constants *K*_d_ for the binding of compounds **20** and **21** to DENV2 protease are measured in the absence or presence of 0.01% TX-100. Binding data were corrected by removing background signal and accounting for inner filter effects using correction factors derived from experiments with NATA.

|  |  | 50 mM Tris, pH 9,<br>1 mM CHAPS<br>His<sub>6</sub>-DENV2 pro<br>RT |  | 50 mM Phosphate, pH 7,<br>1 mM CHAPS<br>Non-tagged DENV2 pro<br>37 degrees |  |
| --- | --- | --- | --- | --- | --- |
|  |  | - TX-100 | + TX-100 | - TX-100 | + TX-100 |
| **20** | $K_d$ | 9.63 ± 1.12 | 6.61 ± 1.88 | 102.27 ± 9.84 | 95.16 ± 21.77 |
| | $IC_{50}$ | 3.01 ± 0.12 | 3.03 ± 0.05 | 49.01 ± 1.86 | 52.51 ± 3.43 |
| **21** | $K_d$ | 3.39 ± 0.08 | 2.00 ± 0.21 | 155.84 ± 15.87 | 119.24 ± 13.03 |
| | $IC_{50}$ | 2.05 ± 0.05 | 2.04 ± 0.03 | 56.25 ± 3.27 | 46.97 ± 2.26 |

Training of machine learning and deep learning models in drug discovery commonly takes place by using datasets of heterogenous inhibition results.^47^ This practice aims to increase the data size in order to improve the model predictivity. However, mixed data are associated with noise and result in uncertainly in the models, setting an upper limit of the performance.^47–49^ It is important thus to be aware of the impact of testing conditions on the data and of the recognition profile of the target protein.

Since compound **2** is prone to degradation by the protease, its stability under different assay conditions was evaluated by LC/MS. For this compound, lower inhibition relative to compounds **3** and **4** in phosphate vs. Tris buffers is not related to degradation of **2** by the protease (cf. Supporting Information, Figure S14). This can be explained by a potential change in the binding mode of the compound. Degradation studies of compound **2** and related *beta*-lactam analogs previously revealed alternate binding modes of the same compound relative to the active site of DENV2pro.^39^

This alternate binding mode can be further supported by the crystal structure of a highly affine boroleucine-based inhibitor bound to ZIKVpro, which shows the feasible fitting of a non-basic side chain in the S1 subpocket.^50^ Using the dipeptide boronic acid compound **1**, we can show for DENVpro that the ligand interactions by the basic recognition residues P1 and P2 experience limited changes with different assay conditions (Table 1). The covalent bond between the boronic acid warhead and the catalytic Ser135 ensures a specific localization of this structural element of the inhibitor. The expected binding mode of **1** (Figure 2B) is based on co-crystallization of a very similar analog, **CN-716**, with ZIKV and WNV proteases,^23,24^ and an NMR study of the binding of a *tert*-butyl tagged derivative to DENV2pro.^25^ **CN-716** is a highly-affine dipeptide boronic acid inhibitor, which has been co-crystallized with WNV and ZIKV proteases (PDB: 5IDK, and 5LC0, respectively). Compound **1** and **CN-716** differ only in the P2 side chain (Arg vs. (4-guanidino)-Phe). As such, binding areas beyond S1 and S2 (prime site, subpocket B, allosteric site) appear to play a role in the sensitivity of dibasic and tribasic substrates as well as inhibitor **2** to buffer composition.

In line with the conformational selection model, and based on the obtained functional evidence, recognition by the protease is proposed to depend on several distinct conformational states. Selected conformations are required to process particular substrates or bind certain inhibitors, and will be populated in solution under specific conditions. In the absence of these conditions, the ligand recognition will not take place to the same extent.

### 3.2. Non-proteolytic recognition in the binding assay

Besides the proteolytic assay, the influence of measurement conditions on the binding of reference inhibitors to DENVpro was assessed in a non-catalytic ligand–protein binding assay. This assay depends on the intrinsic fluorescence of the protein.^13^ Upon binding of a ligand that contains a moiety showing an absorbance spectrum that overlaps with tryptophan emission (330-360 nm), the fluorescence of the protease is quenched in a concentration-dependent manner.^34^ (Behnam, Basché, Klein; Analytical Biochemistry 2023; unformatted draft is provided as review-only supplementary material) To account for optical interference by inner filter effects through absorbance of the excitation or emission wavelengths by the compounds, an experiment is conducted by studying the effect of the compounds on the fluorescence of a solution of *N*-acetyl-L-tryptophanamide (NATA) instead of the protein. Compounds **20-24** (Figure 3A) were used as quenchers, since their N-terminal cap or structure in general shows an absorbance peak between 330 nm and 360 nm. Correction factors derived from experiments with NATA are used to correct the data and confirm specific binding to the target, by showing that the observed quenching of protein fluorescence persists after accounting for any optical interference. Affine non-quenching ligands such as the peptidic inhibitor aprotinin (bovine pancreatic trypsin inhibitor, BPTI) or compounds **25** and **26** (Figure 3A) are used for displacement experiments.

**Figure 3.**
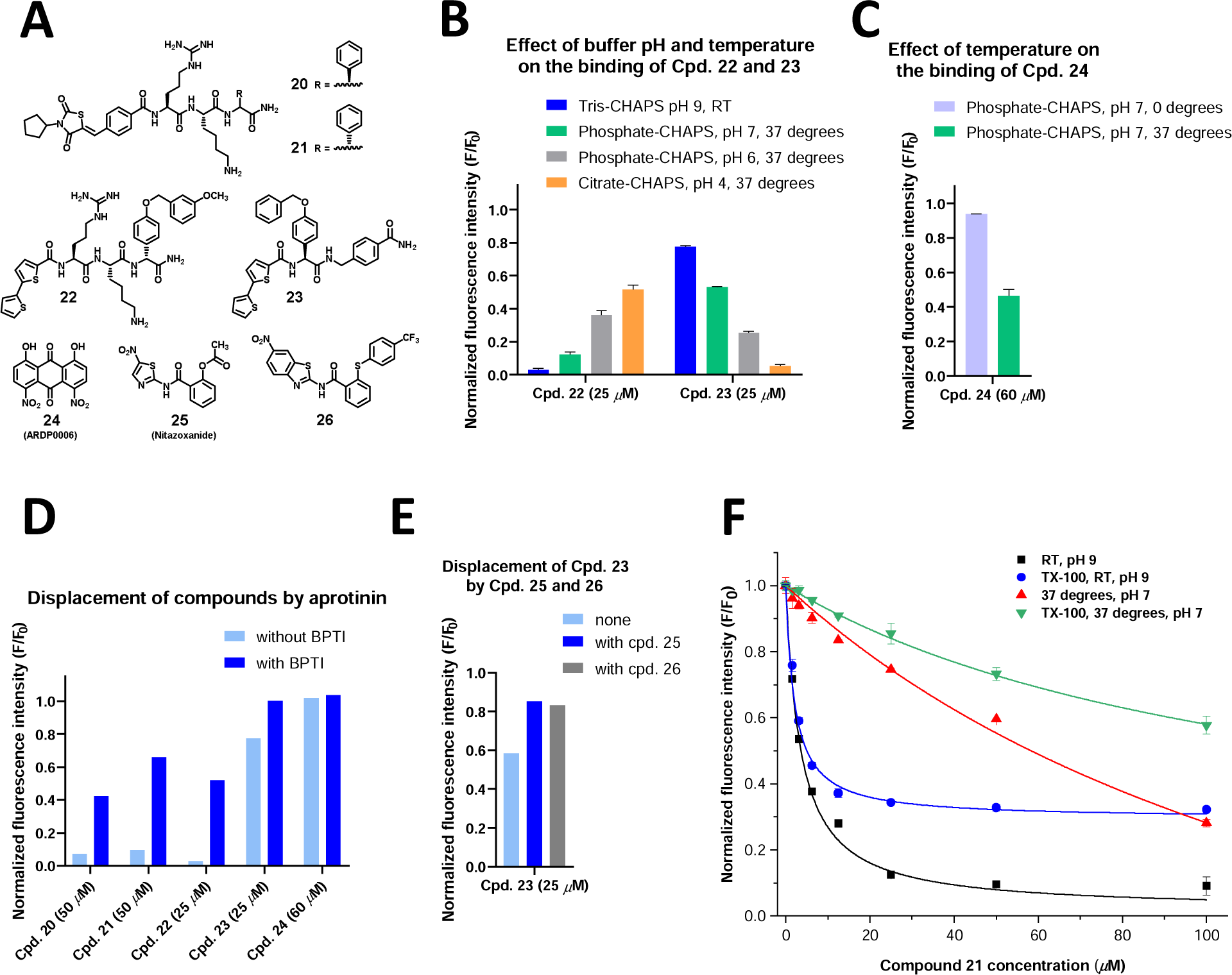
Alternate molecular recognition by DENV2 protease in the binding assay. (**A**) Structure of inhibitors included in the binding assay. (**B**) Effect of buffer pH and temperature on the binding profile of compounds **22** and **23**. (**C**) Effect of temperature on the binding profile of compound **24**. (**D**) Displacement of compounds **20-24** by the highly affine active site inhibitor aprotinin (10 *µ*M) using Tris-CHAPS buffer, pH 9, RT. (**E**) Displacement of compound **23** by two small compounds **25** and **26** (100 *µ*M) using Tris-CHAPS buffer, pH 9, RT. (**F**) Binding of compound **21** to DENV protease in absence or presence of 0.01% TX-100 using two assessment conditions: Tris-CHAPS buffer, pH 9, measurement at RT with His_6_-tagged enzyme; or phosphate-CHAPS buffer, pH 7, measurement at 37 °C with non-tagged enzyme. All plotted binding data were corrected by removing background signal and accounting for inner filter effects using correction factors derived from experiments with NATA.

#### 3.2.1. Feasibility of the binding assay beyond the pH range required for the proteolytic assay

Assessment of ligand binding to the flaviviral protease was not previously conducted under acidic pH. Information about the structure of the protease at pH values below 7 was previously reported by several in-solution techniques.^30,51–53^ Differences in the structure were observed with the decrease in pH, and could be pinpointed to NS2B-C,^28^ His51,^53^ and the overall tertiary packing.^51^ A recent study noticed higher tendency of aggregation of DENVpro at pH 4 following overnight equilibration at room temperature,^52^ while NMR measurement extending over some hours was possible at the same pH in another study.^51^ In comparison to previously published work, the binding assay proceeds with shorter incubation time, of only 1 hr, besides the inclusion of 1 mM CHAPS and the use of higher temperature and lower enzyme concentration.

Temperature and the ionic species of certain buffers at acidic pH reduce the recorded fluorescence for fluorophores.^54–56^ In our hands, this was most obvious with 50 mM acetate buffer, pH 4, 37 °C, where the signal window was almost lost in comparison to the blank, when maintaining the same protein concentration as for other testing conditions such as Tris, pH 9. (cf. Supporting Information, Figure S15) Measurement with a lower signal window was possible in phosphate and citrate buffers, pH 6 and 4, respectively. The signal in this case is comparable to phosphate buffer, pH 7. When needed, the signal window can be increased by higher protein concentrations, but to allow comparison of results between different conditions in this set of experiments, we kept the DENVpro concentration constant at 200 nM.

#### 3.2.2. Assay conditions influence the binding affinity of inhibitors in a non-catalytic binding assay

Similar to the proteolytic assay, the buffer choice and temperature of the measurement influence the extent of ligand–protein binding. Under the tested conditions and using 1% final DMSO concentration (or 2% DMSO for displacement with **25** and **26**), indications of precipitation were absent, where no relevant change in solution turbidity was noted. For the dibasic compounds **20-22**, the binding is reduced by lowering the pH and increasing the temperature (Figure 3B and 3F). The opposite is observed with compounds **23** (Figure 3B) and **24** (ARDP0006) (Figure 3C and Supporting Information, Figure S16). Compound **23** binds to DENVpro in citrate buffer, pH 4, 37 °C with a *K*_d_ of 2.4 ± 0.6 *µ*M. The observed pH-dependent binding of **24** to DENVpro can originate from an overlap between two effects. On the one side, there is the established change in the protein conformation. On the other side, **24** contains ionizable groups (predicted p*K*_a_ 3.5) and hence the distribution of species in solution is expected to vary. As previously discussed, the pH of phosphate buffer has the least change with temperature. This property allows to observe the specific effect of temperature on the binding of **24** to DENVpro (Figure 3C). Inhibitors **20-24** are displaced by aprotinin (BPTI), resulting in restoration of protein fluorescence (Figure 3D and Supporting Information, Figures S17 and S18), and **23** is also displaced by compounds **25** and **26** (Figure 3E).

NMR studies reported an influence of the pH on the distribution of open and closed conformations in solution, where DENVpro predominantly populates the open conformation at higher pH and the closed conformation at lower pH.^28,53^ These pH-related conformational changes were linked by mutagenesis studies to His51.^53^ Recent work by Basché, Schirmeister and colleagues^57^ showed that a dibasic active site inhibitor, **20**, shifts the conformational equilibrium in solution towards the closed conformation. However, an allosteric inhibitor results instead in a shift from the closed towards the open conformation. To show the effect of the allosteric inhibitor in Tris-CHAPS buffer, pH 9, the closed conformation was first stabilized by the addition of the dibasic active site inhibitor. The tendency of some inhibitors to bind or stabilize different conformations may explain the distinct binding profiles of inhibitors **22** vs. **23**. The closed conformation stabilized through the addition of a dibasic or tribasic substrate or inhibitor may differ in the space distribution available for further binding in comparison to that obtained by lower pH. In this case, the first is a ligand–enzyme complex, while the second is an unbound enzyme. In the absence of ligand-related justification, the distinct influence of measurement conditions on binding parameters supports a molecular recognition based primarily on conformational selection.

The effect of temperature alone independent from the pH can be observed for compound **24**, where in the same buffer, phosphate-CHAPS pH 7, incubation at 0 °C shows no specific binding to the target in comparison to 37 °C (Figure 3C). For compounds **22** and **23**, the binding was enhanced in phosphate-CHAPS pH 7 at 37 °C in comparison to 0 °C. The impact of temperature on recognition is more challenging to interpret as, to the best of our knowledge, no structural studies evaluated the influence of this factor on the protease structure. Only one report noted higher dispersion of NMR signals of the protease at 37 °C vs. 25 °C.^29^ Structural studies in solution can be conducted at higher temperatures such as 25 °C or 37 °C in comparison to lower temperatures commonly used for X-ray crystallization. Therefore, we consider it likely that temperature may be connected to the distinct conformations reported for DENVpro by NMR (PDB: 2M9P and 2M9Q)^27^ or to other higher-energy conformation.^58^ Compound **24** was shown to inhibit an intramolecular cleavage performed by the dengue protease, consequently preventing the NS2B–NS3 autocleavage.^59^

#### 3.2.3. Non-ionic detergents have different effects in proteolytic vs. binding assays

The non-ionic detergent Triton X-100 (TX-100) was previously reported to interfere with inhibitory activity of certain compounds.^14,15^ This is also observed for the binding of compounds **20-23** and with another non-ionic detergent, Brij 58 (cf. Supporting Information, Figure S19). However, assessment of inhibition by compounds **20** and **21** in the proteolytic assay, in the absence and presence of TX-100, does not show a similar effect as observed in the binding assay (Figure 3F and Table 2). TX-100 was previously reported to interfere with the interaction of the NS2B N-terminus with NS3pro.^45^ Therefore, one possible explanation is that the binding assay can detect binding at other sites than the catalytic active site, and that the assessed compounds have more than one binding position, similar to an interpretation previously suggested for compound **2**.^3,39^ Alternatively, the tribasic AMC substrate may exert distinct influence on the conformational equilibrium in comparison to the used ligand. Differences between *K*_d_ and IC_50_ values obtained from binding and proteolytic assays were also observed for other compounds.^34,60^ (Behnam, Basché, Klein; Analytical Biochemistry 2023; manuscript accepted for publication; unformatted draft is provided as review-only supplementary material) In the binding assay, the presence of TX-100 lowers the maximum binding of compounds **20** and **21** to the protease, and slightly enhances the *K*_d_, suggesting thus an uncompetitive mode of inhibition.

## 4. CONCLUSION

The present manuscript is the first step towards functionally demonstrating alternate recognition of DENVpro, based on a mechanism that relies primarily on conformational selection. Our results show the impact of testing conditions in proteolytic and binding assays for DENVpro. The influence of certain factors on proteolytic processing is substrate-specific, and is distinct from factors that affect the catalytic competence of the core active site, which depends upon the configuration of the catalytic triad and the extent of serine electrophilicity. Recognition of structurally distinct ligands depends to a variable degree on the choice of assay conditions, altering thus the relative ranking of hits. This explains the performance limitations of artificial intelligence models relying on heterogenous inhibition data. Understanding the recognition profile of the target protein can provide an opportunity to overcome this challenge. From the results of our cellular assay system DENV2proHeLa, it becomes evident that adjusting testing systems based on the conditions for optimum processing for a particular substrate may not correlate with cellular assays. This highlights the importance of hit classification based on cellular assay systems, in which the target is present under physiological conditions and ideally in its native microenvironment.

Furthermore, the binding assay identifies distinct effects for certain classes of ligands in comparison to the proteolytic assay. The choice of the most relevant assay conditions for hit identification and optimization would require large-scale correlations. However, we do not exclude potential complementarity between non-similar experimental procedures, based on the recognition of selected scaffolds by different conformations in solution. Additionally, while the simplified assays on the isolated protease are valuable tools, it is possible that they are unable to capture the full conformational landscape of the protease in the cellular systems. The unique functional insights obtained in this work are based on results and observations accumulating from multidisciplinary approaches and efforts to understand the binding of inhibitors, and ligands in general, to flaviviral proteases. These insights provide supportive evidence to the conformational selection model that can be added to experimental findings obtained by other groups in the field. Altered function and molecular recognition by DENVpro depending on the surrounding environment can be explained by the conformational plasticity noted by structural studies. It is likely that some of these states have a different function in the viral replication process and that certain states are more relevant for antiviral strategies than others. Notably, this effect was shown for the development and efficacy of vaccines based on the SARS-CoV-2 spike protein, where a particular conformation is linked to the generation of neutralizing antibodies.^61^

## AUTHOR CONTRIBUTIONS

M.B. and C.K. designed the study. M.B. performed the experimental work. M.B. and C.K. analyzed the results. M.B. and C.K. wrote the manuscript.

## ACKNOWLEDGMENTS

We appreciate the synthesis of inhibitor **1** by Dr. Mascha Dieckmann at the Chemical Biology Core Facility, EMBL, Heidelberg. We thank Dr. Stefan Hinkes for synthetic optimization of the AMC substrate, Dr. Nikos Kühl, Dr. Christian Gege and Yuxing Deng for provided inhibitors, and Katharina Eckstein for assistance in protein expression. We acknowledge valuable comments on the manuscript from Dr. Nikos Kühl, Dr. Cheng Zhang, Johannes Lang and Leah Glanzmann. Furthermore, we thank Heiko Rudy for ESI-MS and LC/MS measurements, Tobias Timmermann for NMR measurements, and Natascha Stefan for technical assistance. C.K. acknowledges generous financial support from the Volkswagen-Stiftung for the project “Preclinical development of antiviral protease inhibitors targeting flavi- and coronaviruses” (9A836). Multiple inhibitors reported in this work were obtained within a project supported by the Deutsche Forschungsgemeinschaft under grant No. 1356/3. The synthesis of inhibitor **1** was originally carried out in a project which received financial support by the German Center for Infection Research (DZIF, TTU 01.911).

## SUPPORTING INFORMATION

Supporting information is provided including abbreviations, analytical characterization of synthesized substrates and inhibitors, kinetic characterization of substrates, assessment of compound degradation by LC/MS, in addition to supplementary figures and tables for the proteolytic and non-proteolytic binding assays.

