## Supplemental Information for "Alternate recognition by dengue protease: Proteolytic and binding assays provide functional evidence beyond an induced-fit"

### Supporting information

**Abbreviations**

EG, ethylene glycol

GL, glycerol

FRET, Förster or fluorescence resonance energy transfer

AMC, 7-amino-4-methylcoumarin

*p*NA, *para*-nitroaniline

RT, room temperature

NATA, *N*-acetyl-L-tryptophanamide

**Analytical characterization of substrates**

| **Cpd.** | **Molecular formula** | **Calc. Mass $[M+H]^+$** | **Found mass $[M+H]^+$** | **Yield (%)** | **Purity by HPLC at 254 nm (%)** |
|---|---|---|---|---|---|
| Substrate **6** | $C_{43}H_{68}N_{16}O_{11}$ | 985.5326 | 985.5296 | 20.1 | 100 |
| Substrate **7** | $C_{62}H_{90}N_{18}O_{20}$ | 1407.6652 | 1407.6598 | 27.2 | 91 |
| Substrate **8** | $C_{52}H_{80}N_{22}O_{14}$ | 1237.6297 | 1237.6296 | 33.5 | 97 |
| Substrate **9** | $C_{59}H_{85}N_{19}O_{13}$ | 1268.6647 | 1268.6606 | 28.9 | 96 |
| Substrate **10** | $C_{45}H_{69}N_{15}O_{13}$ | 1028.5272 | 1028.5263 | 32.7 | 100 |
| Substrate **11** | $C_{43}H_{68}N_{18}O_{12}$ | 1029.5337 | 1029.5322 | 29.8 | 94 |
| Substrate **12** | $C_{48}H_{71}N_{17}O_{15}$ | 1126.5388 | 1126.5405 | 4.4 | 100 |
| Substrate **13** | $C_{56}H_{81}N_{17}O_{12}$ | 1184.6323 | 1184.6309 | 21.8 | 91 |
| Substrate **14** | $C_{58}H_{79}N_{17}O_{16}$ | 1270.5963 | 1270.5936 | 15.9 | 96 |
| Substrate **15** | $C_{53}H_{74}N_{16}O_{13}$ | 1143.5694 | 1143.5721 | 15.7 | 100 |
| Substrate **16** | $C_{30}H_{46}N_{10}O_6$ | 643.3675 | 643.3676 | 66.7* | 97 |
| Substrate **17** | $C_{41}H_{60}N_{12}O_7$ | 833.4781 | 833.4764 | 65.9* | 100 |
| Substrate **19** | $C_{26}H_{43}N_{11}O_6$ | 606.3471 | 606.3480 | 70.8* | 100 |
| | | **Calc. Mass $[M-H]^-$** | **Found mass $[M-H]^-$** | | |
| Intermediate **Ia** | $C_{27}H_{43}N_5O_7S$ | 580.2810 | 580.2833 | | |
| Intermediate **Ib** | $C_{43}H_{65}N_7O_{10}S$ | 870.4441 | 870.4434 | | |

* Yields for the last coupling step in solution.

**Table S1** Characterization and purity assessment of synthesized substrates

**Analytical characterization of inhibitors**

Compound **1**: $^1$H NMR ($D_2O$) δ 7.78 (d, *J* = 8 Hz, 2H), 7.63 (m, 1H), 7.54 (m, 2H), 4.66 (m, 1H), 3.25 (s, 2H), 3.16 (s, 2H), 2.65 (m, 1H), 1.94 (m, 2H), 1.75 (m, 2H), 1.57 (m, 4H). UPLC-MS, m/z $[M-H_2O+H]^+$ for $C_{18}H_{30}BN_8O_4$: calculated 417, found: 417.

Compound **2**: HRMS (ESI), m/z $[M + H]^+$ for $C_{34}H_{45}N_8O_5$: calculated, 645.3507; found, 645.3493.

Compound **20**: $^1$H NMR (500 MHz, $D_2O$) δ 7.81 – 7.72 (m, 3H), 7.60 – 7.52 (m, 2H), 7.42 (s, 5H), 5.44 (s, 1H), 4.70 (d, *J* = 7.0 Hz, 1H), 4.47 (td, *J* = 6.7, 3.2 Hz, 2H), 3.19 – 3.10 (m, 2H), 2.99 (t, *J* = 7.6 Hz, 2H), 1.97 – 1.57 (m, 18H), 1.54 – 1.42 (m, 2H).

$^1$H NMR (500 MHz, DMSO-$d_6$) δ 8.65 (d, *J* = 7.7 Hz, 1H), 8.28 (d, *J* = 7.8 Hz, 1H), 8.19 (d, *J* = 8.1 Hz, 1H), 7.99 (d, *J* = 8.6 Hz, 2H), 7.94 (s, 1H), 7.75 – 7.66 (m, 5H), 7.56 (t, *J* = 5.7 Hz, 1H), 7.40 (d, *J* = 8.4 Hz, 2H), 7.36 – 7.25 (m, 4H), 7.19 (s, 1H), 5.33 (d, *J* = 7.7 Hz, 1H), 4.68 (p, *J* = 8.5 Hz, 1H), 4.51 – 4.42 (m, 1H), 4.40 – 4.33 (m, 1H), 3.10 (q, *J* = 7.0 Hz, 2H), 2.74 (d, *J* = 5.6 Hz, 2H), 2.06 – 1.45 (m, 17H), 1.40 – 1.29 (m, 2H), 1.22 (s, 1H). HRMS (ESI), m/z $[M + H]^+$ for $C_{36}H_{48}N_9O_6S$: calculated, 734.3443; found, 734.3446.

Compound **21**: $^1$H NMR (500 MHz, $D_2O$) δ 7.88 (d, *J* = 2.1 Hz, 2H), 7.87 (s, 1H), 7.68 (d, *J* = 8.6 Hz, 2H), 7.50 – 7.43 (m, 5H), 5.46 (s, 1H), 4.75 – 4.73 (m, 1H), 4.53 (t, *J* = 7.2 Hz, 1H), 4.42 (dd, *J* = 8.5, 6.2 Hz, 1H), 3.23 (t, *J* = 6.9 Hz, 2H), 2.92 (t, *J* = 7.9 Hz, 2H), 2.03 – 1.62 (m, 18H), 1.44 – 1.34 (m, 2H).

$^1$H NMR (500 MHz, DMSO-$d_6$) δ 8.66 (d, *J* = 7.6 Hz, 1H), 8.51 (d, *J* = 8.1 Hz, 1H), 8.18 (d, *J* = 8.1 Hz, 1H), 8.04 – 7.99 (m, 2H), 7.96 (s, 1H), 7.76 – 7.64 (m, 6H), 7.59 (d, *J* = 6.7 Hz, 1H), 7.43 – 7.39 (m, 2H), 7.36 – 7.32 (m, 2H), 7.30 – 7.26 (m, 1H), 7.24 (s, 1H), 5.39 (d, *J* = 8.0 Hz, 1H), 4.70 (d, *J* = 8.3 Hz, 1H), 4.52 – 4.37 (m, 2H), 3.31 (s, 1H), 3.16 – 3.08 (m, 2H), 2.70 (s, 2H), 2.04

– 1.78 (m, 8H), 1.75 – 1.45 (m, 10H), 1.33 – 1.22 (m, 2H). HRMS (ESI), m/z $[M + H]^+$ for $C_{36}H_{48}N_9O_6S$: calculated, 734.3443; found, 734.3441.

Compound **22**: HRMS (ESI), m/z $[M + H]^+$ for $C_{37}H_{47}N_8O_6S_2$: calculated, 763.3054; found, 763.3037.

Compound **23**: HRMS (ESI), m/z $[M + H]^+$ for $C_{32}H_{28}N_3O_4S_2$: calculated, 582.1516; found, 582.1514.

**Supplementary figures for the proteolytic assay**

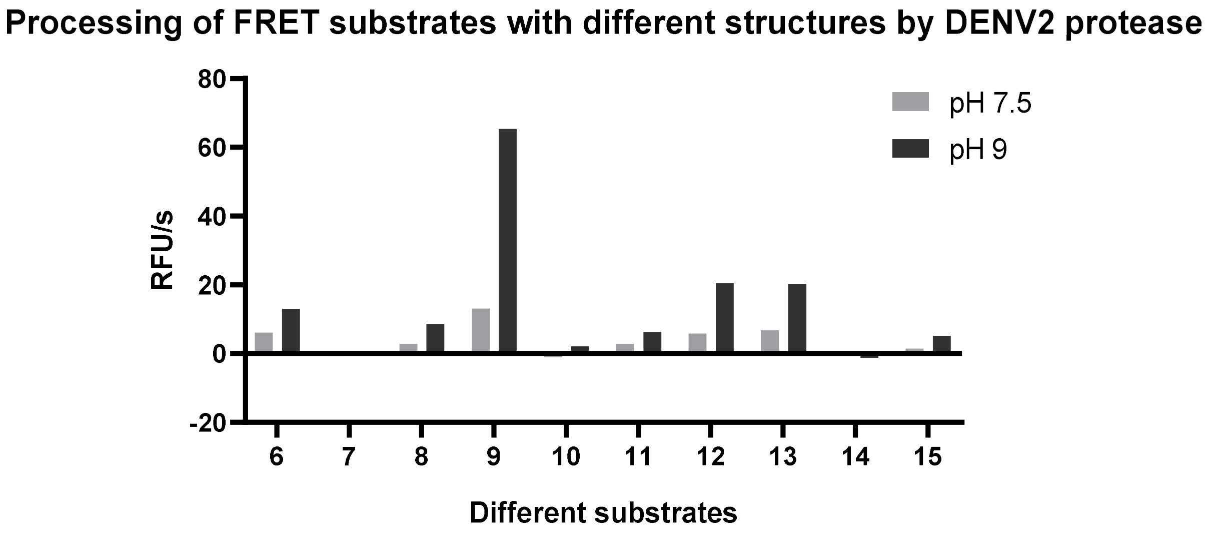

**Figure S1** Proteolytic processing of FRET substrates with different structures by DENV2 protease. Activity is indicated as slope per second of relative fluorescence units (RFU/s). $His_6$-tagged linked construct is used. All FRET substrates are cleaved preferentially in Tris buffer pH 9 compared to pH 7.5 (the buffer is 50 mM Tris, without additives). The highest signal is observed for substrate **9**, which is based on the NS2A/2B cleavage site.

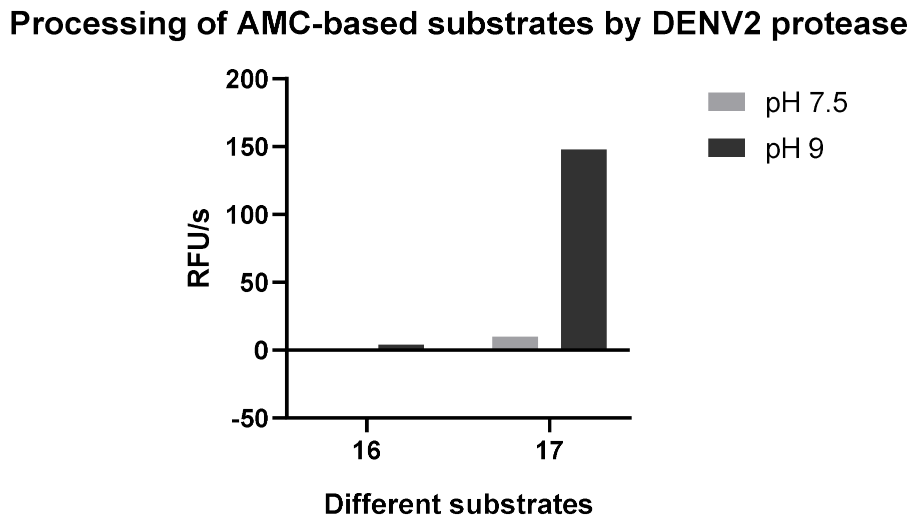

**Figure S2** Proteolytic processing of AMC-based substrates by DENV2 protease. Cleavage is higher in Tris buffer pH 9 relative to pH 7.5, and for the tribasic substrate **17** more than the dibasic substrate **16**. Activity is indicated as slope per second of relative fluorescence units (RFU/s). $His_6$-tagged linked construct is used. The buffer is 50 mM Tris without additives.

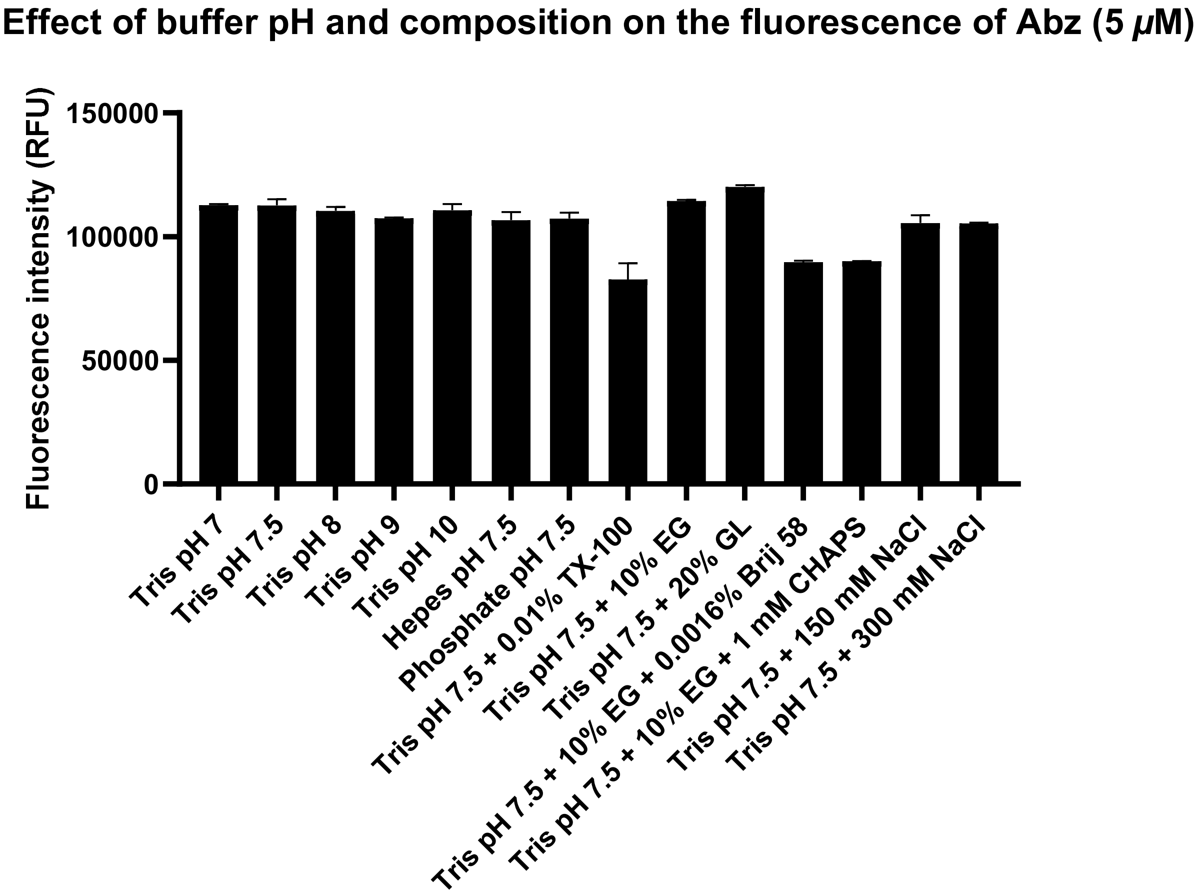

**Figure S3** No significant effect of the buffer pH and composition can be seen on the fluorescence of Abz (5 $\mu$M).

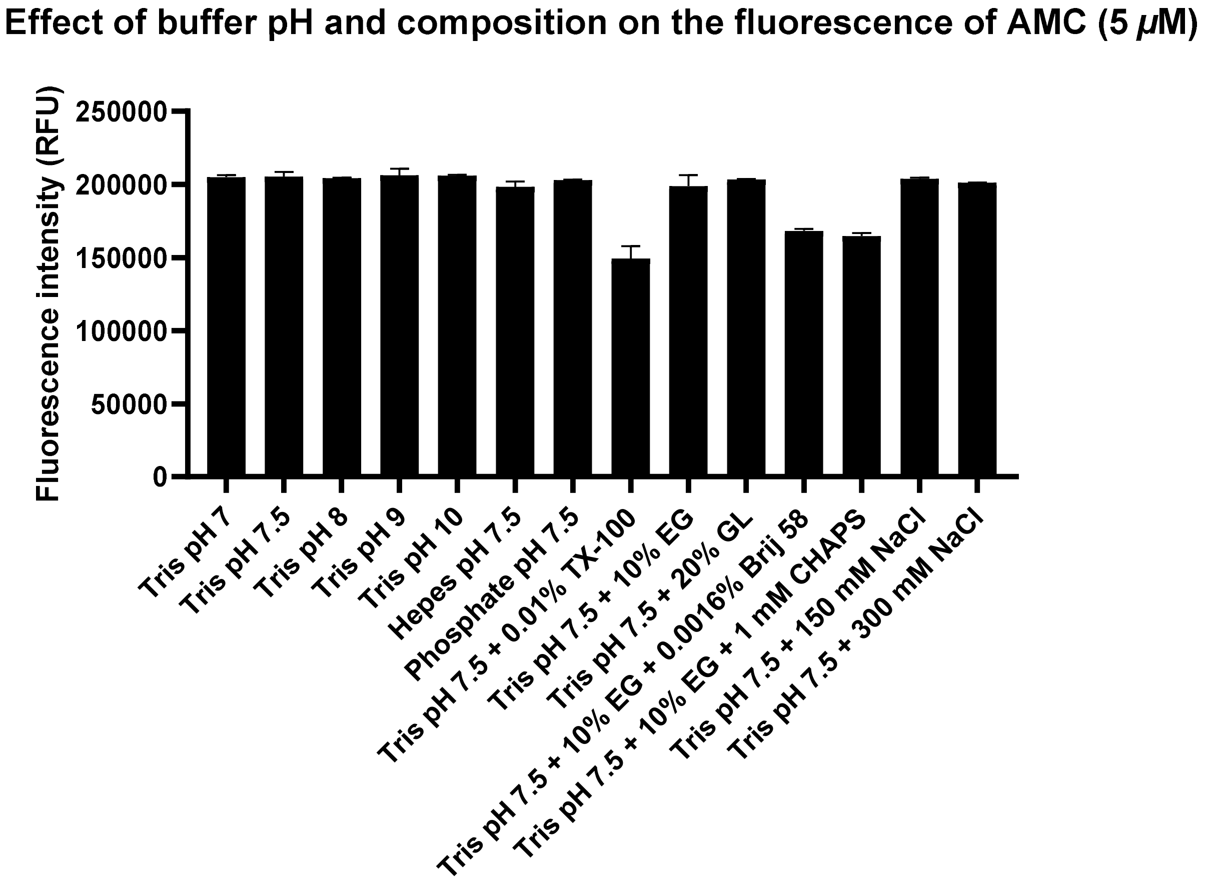

**Figure S4** No significant effect of the buffer pH and composition can be seen on the fluorescence of AMC (5 $\mu$M).

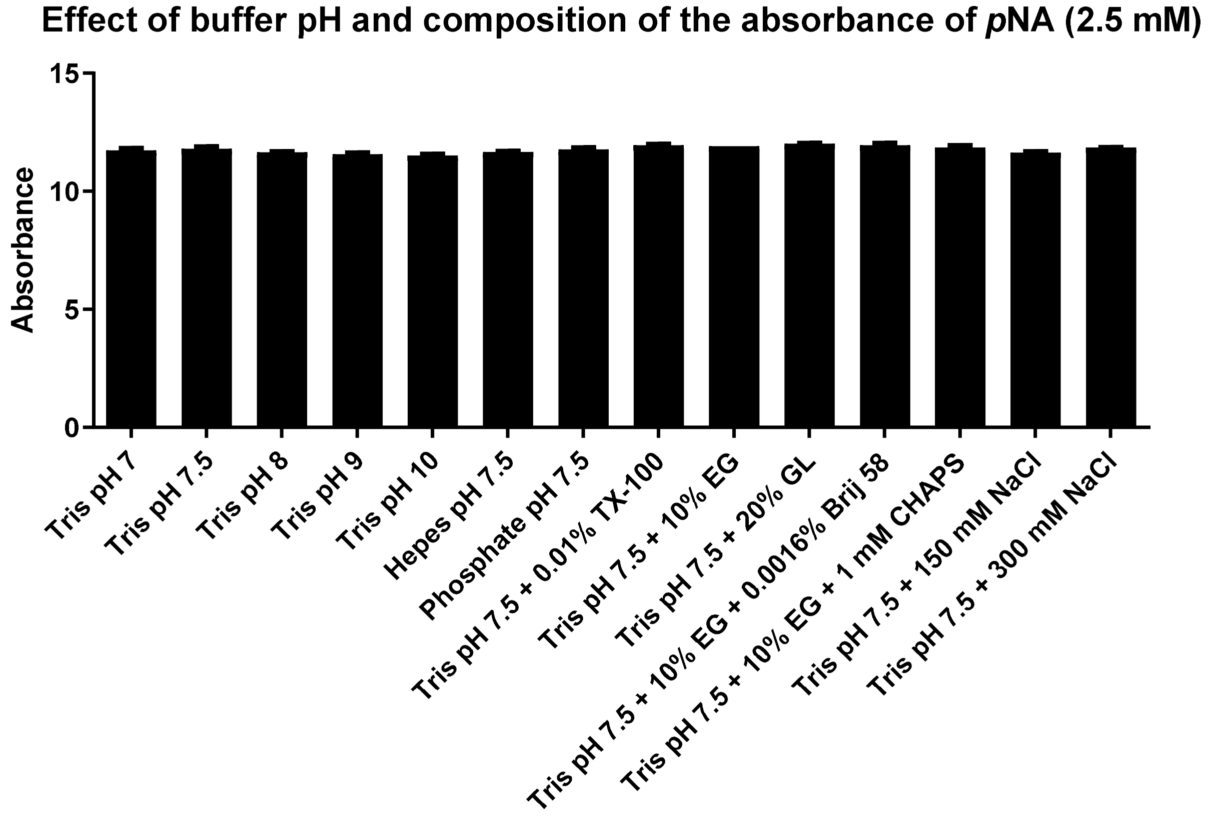

**Figure S5** No significant effect of the buffer pH and composition can be seen on the absorbance of *p*NA (2.5 mM).

**pH profile in Tris buffer for the AMC-based substrate 16 on DENV2 protease**

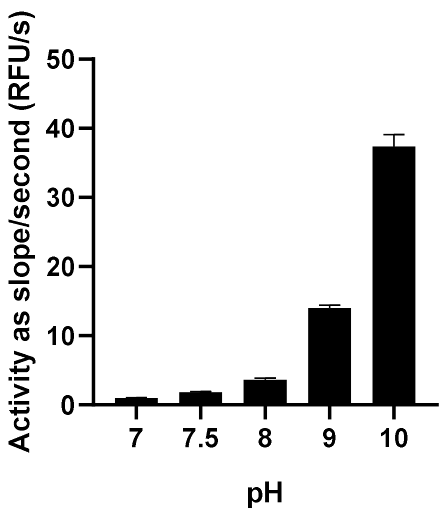

**Figure S6** pH profile of the AMC-based substrate **16** on DENV2 protease. All buffers are prepared as 50 mM without additives. $His_6$-tagged linked construct of DENV2pro is used.

**pH profile in Tris buffer for the *p*NA-based substrate 19 on DENV2 protease**

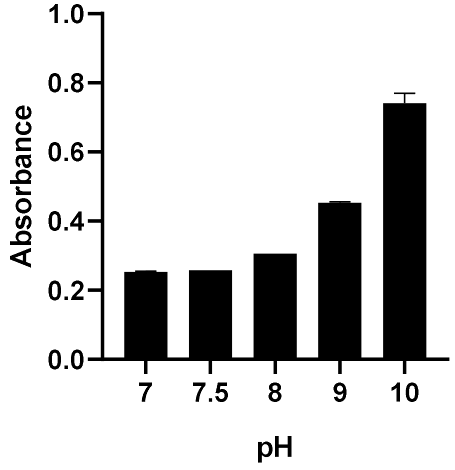

**Figure S7** pH profile of the *p*NA-based substrate **19** on DENV2 protease. All buffers are prepared as 50 mM without additives. $His_6$-tagged linked construct of DENV2pro is used. The pH profile is less steep between pH 7 and pH 10 in comparison to substrates **9** or **17** (refer to Figure **2** in main manuscript).

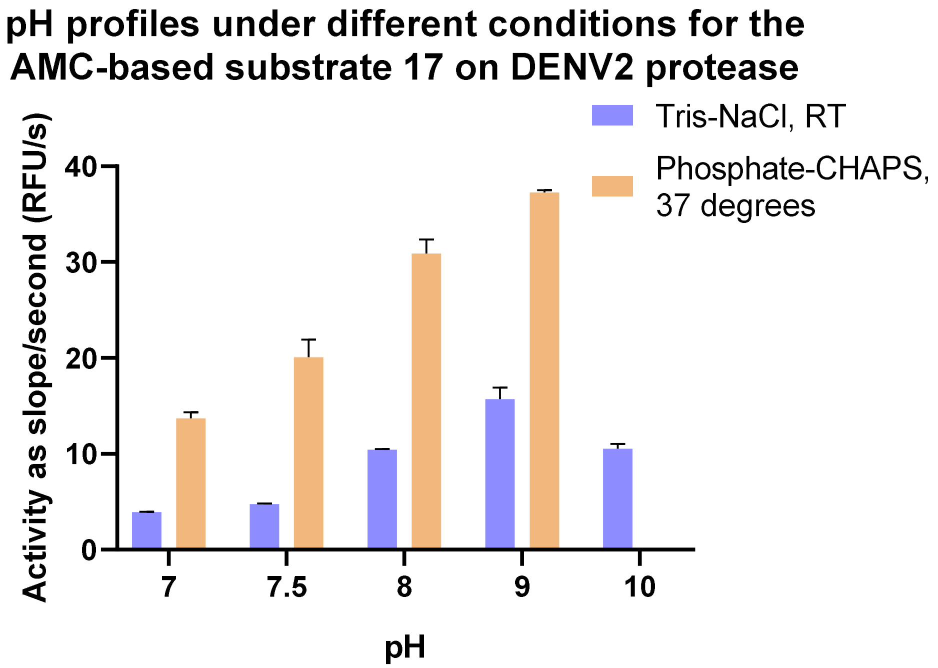

**Figure S8** pH profiles of the AMC-based substrate **17** on DENV2 protease. The pH profile for the same substrate, **17**, varies under different measurement conditions. For the phosphate-CHAPS buffer, pH 10 was not assessed (refer to Figure **2** in main manuscript). All buffers are prepared as 50 mM without additives. $His_6$-tagged linked construct of DENV2pro is used.

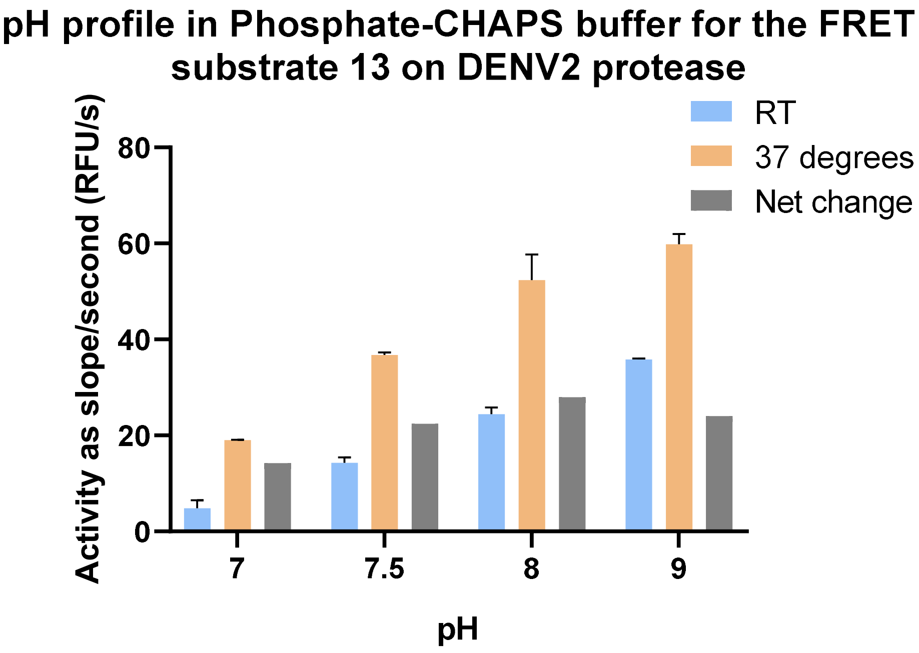

**Figure S9** Plot for the pH profiles for the proteolytic processing of the FRET substrate **13** by DENV2 protease. All buffers are prepared as 50 mM without additives. The non-tagged linked construct of DENV2pro is used. The net change in the readings between the measurements at 37 degrees and RT is highest at pH 8.

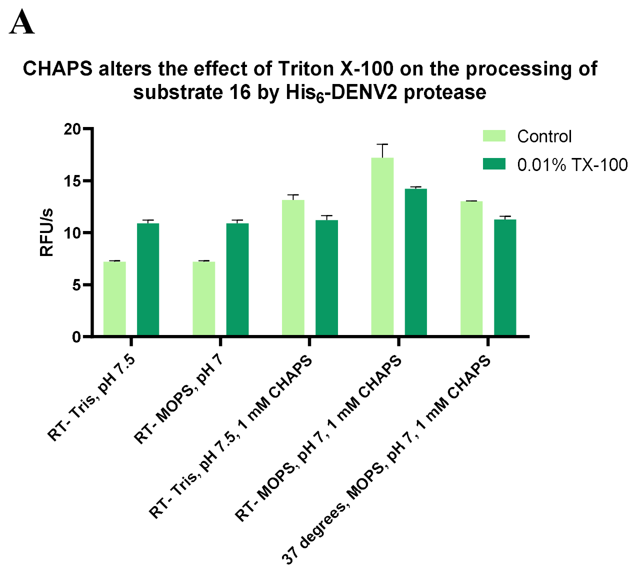

**Figure S10** CHAPS alters the effect of Triton X-100 on the proteolytic processing of substrate **16** by DENV2 protease. The presented results were obtained using the $His_6$-tagged linked construct in (A) or the non-tagged linked construct in (B). The results show that the observed effect is independent of the $His_6$-tag.

**Buffer-dependent temperature effect for processing the FRET substrate 15 by DENV2 protease at pH7**

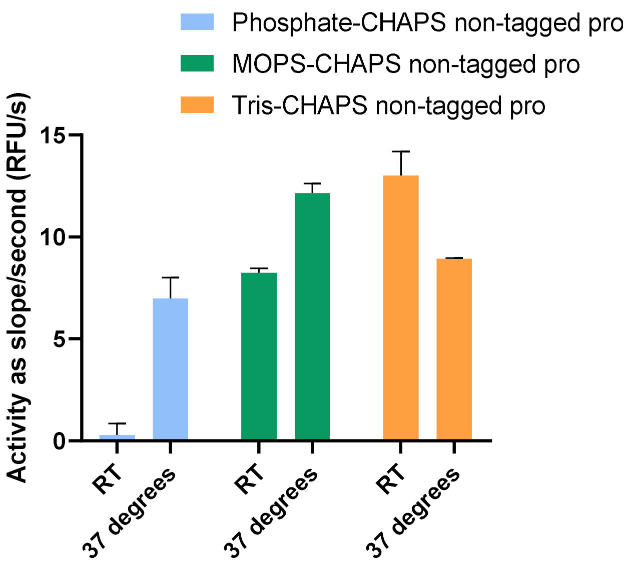

**Figure S11** Temperature effect in different buffers for the proteolytic processing of the FRET substrate **15** by DENV2 protease at pH 7. All buffers are prepared as 50 mM, and CHAPS concentration is 1 mM. The non-tagged construct of DENV2pro is used.

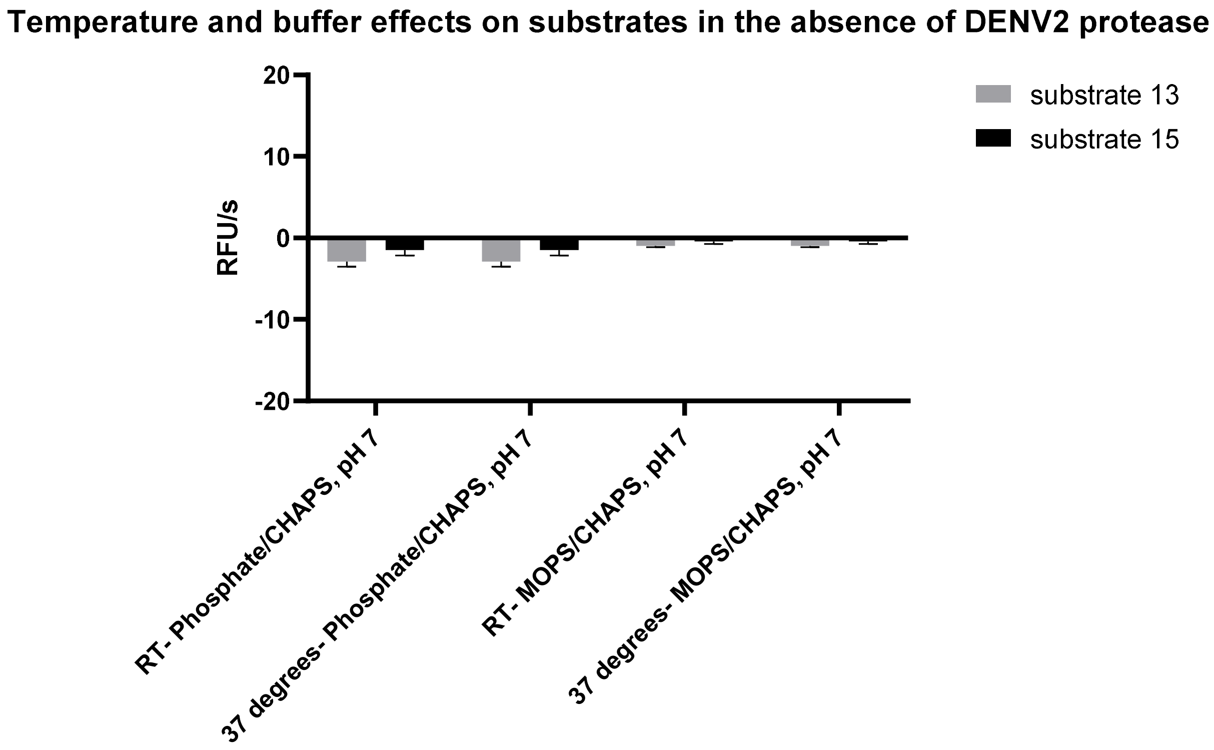

**Figure S12** Blank readings in the absence of the enzyme. Buffers are prepared as 50 mM and CHAPS has a concentration of 1 mM.

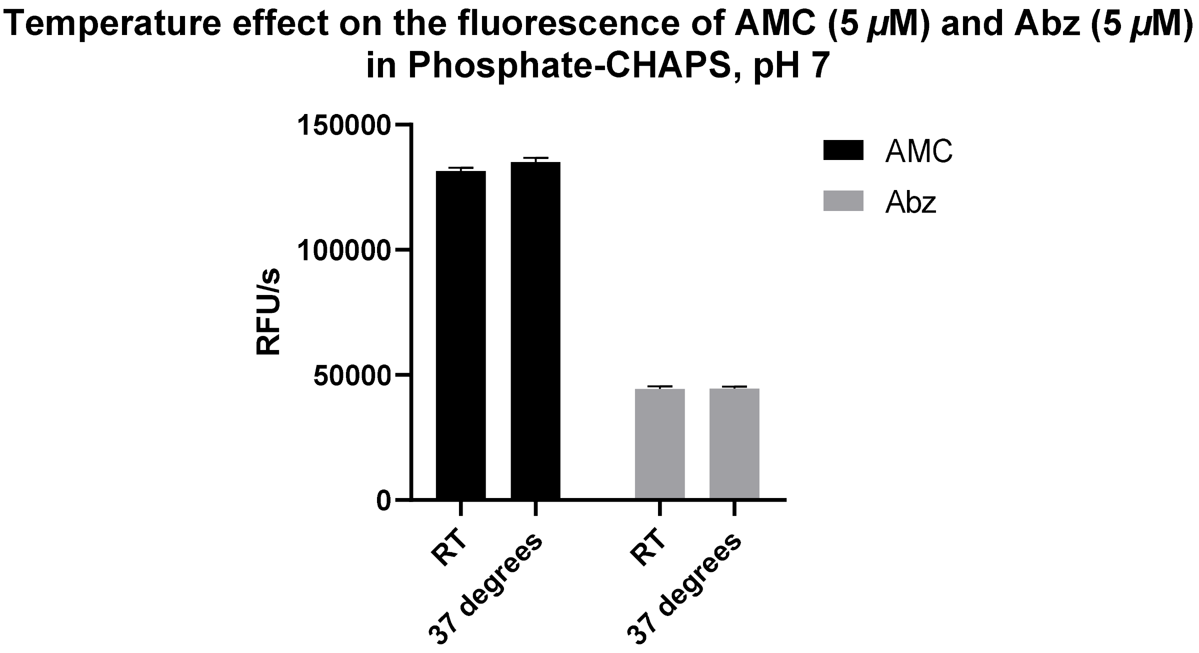

**Figure S13** No effect for the change in temperature is detected on the fluorescence of AMC or Abz (5 $\mu$M).

#### Kinetic characterization of substrates

| | | |
|---|---|---|
| **Structure** | **FRET substrate 9**<br>2-Abz-LKRRSWP-(3-$NO_2$)Y-$NH_2$ | **AMC-based substrate 17**<br>Bz-nKRR-AMC |
| **Buffer 1:** 50 mM Tris, pH 9, 10% EG, 1 mM CHAPS, RT | | |
| **$K_m$ ($\mu$M)** | 14.40 ± 2.925 | 15.86 ± 1.19 |
| **$V_{max}$ ($\mu$M/s)** | 0.03155 ± 0.002088 | 0.0152 ± 0.0003833 |
| **$K_{cat}$ ($s^{-1}$)** | 0.3155 ± 0.02088 | 0.3040 ± 0.007667 |
| **$K_{cat}$ / $K_m$ ($M^{-1}s^{-1}$)** | 21909.7222 | 19167.7175 |
| **Buffer 2:** 50 mM Tris, pH 7.5, 10% EG, 1 mM CHAPS, RT | | |
| **$K_m$ ($\mu$M)** | 138.9 ± 23.78 | 90.36 ± 4.573 |
| **$V_{max}$ ($\mu$M/s)** | 0.01987 ± 0.001825 | 0.0108265 ± 0.0001952 |
| **$K_{cat}$ ($s^{-1}$)** | 0.1987 ± 0.01825 | 0.08265 ± 0.001952 |
| **$K_{cat}$ / $K_m$ ($M^{-1}s^{-1}$)** | 1430.5256 | 914.6746 |
| **Buffer 3:** 50 mM Phosphate, 300 mM NaCl, pH 7.5, 10% EG, 1 mM CHAPS, RT | | |
| **$K_m$ ($\mu$M)** | 348.7 ± 28.69 | 149.0 ± 13.75 |
| **$V_{max}$ ($\mu$M/s)** | 0.01122 ± 0.0006491 | 0.002021 ± 0.0001023 |
| **$K_{cat}$ ($s^{-1}$)** | 0.1122 ± 0.006491 | 0.02021 ± 0.001023 |
| **$K_{cat}$ / $K_m$ ($M^{-1}s^{-1}$)** | 321.7666 | 135.6376 |
| **Buffer 3:** 50 mM Phosphate, 300 mM NaCl, pH 7.5, 10% EG, 1 mM CHAPS, RT<br>Enzyme: Non-tagged DENV2pro | | |
| **$K_m$ ($\mu$M)** | 951 ± 90.92 | 262.4 ± 17.27 |
| **$V_{max}$ ($\mu$M/s)** | 0.03309 ± 0.002703 | 0.002768 ± 0.0001187 |
| **$K_{cat}$ ($s^{-1}$)** | 0.3309 ± 0.02703 | 0.02768 ± 0.001187 |
| **$K_{cat}$ / $K_m$ ($M^{-1}s^{-1}$)** | 347.9495 | 105.4878 |
| **Buffer 4**: 50 mM Phosphate, pH 7, 1 mM CHAPS, 37 °C | | |
| **$K_m$ ($\mu$M)** | n.d. | 75.52 ± 4.925 |
| **$V_{max}$ ($\mu$M/s)** | n.d. | 0.001433 ± 0.0000423 |
| **$K_{cat}$ ($s^{-1}$)** | n.d. | 0.01433 ± 0.000423 |
| **$K_{cat}$ / $K_m$ ($M^{-1}s^{-1}$)** | n.d. | 189.7511 |
| **Structure** | **FRET substrate 13**<br>2-Abz-VKKR-AAW-(3-$NO_2$)Y | |
| **Buffer 4**: 50 mM Phosphate, pH 7, 1 mM CHAPS, 37 °C | | |
| **$K_m$ ($\mu$M)** | 543.5 ± 134.1 | |
| **$V_{max}$ ($\mu$M/s)** | 0.04808 ± 0.009095 | |
| **$K_{cat}$ ($s^{-1}$)** | 0.09615 ± 0.01819 | |
| **$K_{cat}$ / $K_m$ ($M^{-1}s^{-1}$)** | 176.9089 | |
| **Structure** | **AMC-based substrate 16**<br>(*t*-Bu)acetic acid-GRR/AMC | |
| **Buffer 2:** 50 mM Tris, pH 7.5, 10% EG, 1 mM CHAPS | | |
| **$K_m$ ($\mu$M)** | 182.5 ± 7.87 | |
| **$V_{max}$ ($\mu$M/s)** | 0.001676 ± 0.0000423 | |

| | |
|---|---|
| $\boldsymbol{K}_{\mathbf{cat}}$ **(s$^{-1}$)** | 0.003353 ± 0.0000845 |
| $\boldsymbol{K}_{\mathbf{cat}}$ **/** $\boldsymbol{K}_{\mathbf{m}}$ **(M$^{-1}$s$^{-1}$)** | 18.3726 |
| **Buffer 2:** 50 mM Tris, pH 7.5, 10% EG, 1 mM CHAPS<br>Enzyme: Non-tagged DENV2pro | |
| $\boldsymbol{K}_{\mathbf{m}}$ **($\mu$M)** | 220.8 ± 8.44 |
| $\boldsymbol{V}_{\mathbf{max}}$ **($\mu$M/s)** | 0.001940 ± 0.000046 |
| $\boldsymbol{K}_{\mathbf{cat}}$ **(s$^{-1}$)** | 0.003880 ± 0.000092 |
| $\boldsymbol{K}_{\mathbf{cat}}$ **/** $\boldsymbol{K}_{\mathbf{m}}$ **(M$^{-1}$s$^{-1}$)** | 17.5724 |

**Table S2** Kinetic parameters for the proteolytic processing of substrates by DENV2 protease under different buffers. $His_6$-tagged linked construct of DENV2pro was used, unless otherwise indicated. All measurements were performed using an enzyme concentration of 100 nM, except for the following: buffer 1 for the AMC-based substrate **17**, enzyme concentration 50 nM: buffer **4** for the FRET substrate **13** and buffer **2** for the AMC-based substrate **16**, enzyme concentration 500 nM. The term n.d. stands for not-determined. For substrate **9**, substrate inhibition was observed in the Michaelis-Menten curve with buffer 1 at concentrations starting from 200 $\mu$M, while the same was not seen with other buffers. RT stands for room temperature.

**Assessment of compound degradation by LC/MS**

**14a** Buffer: 50 mM Tris, pH 9, 1 mM CHAPS, RT

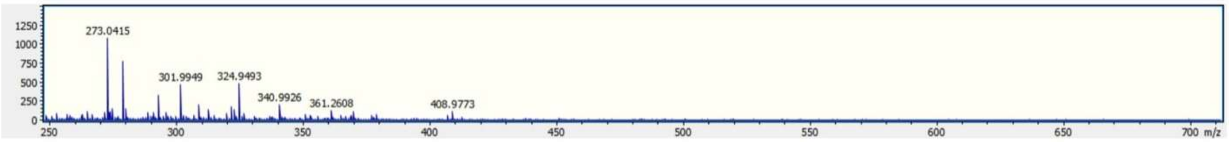

**14b** Buffer: 50 mM Tris, pH 7.5, 1 mM CHAPS, RT

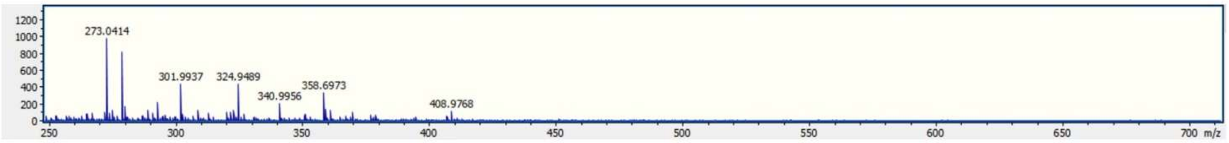

**14c** 50 mM Phosphate, pH 7.5, 300 mM NaCl, 1 mM CHAPS, RT

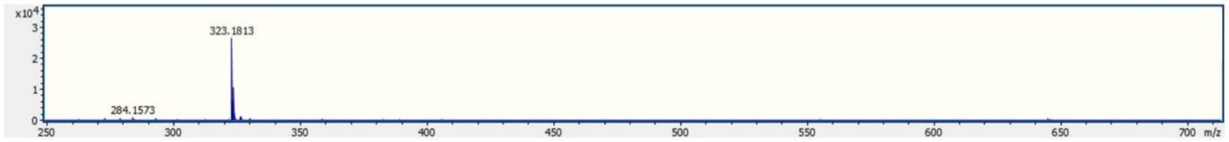

**14d** 50 mM Phosphate, pH 7, 1 mM CHAPS, 37 °C

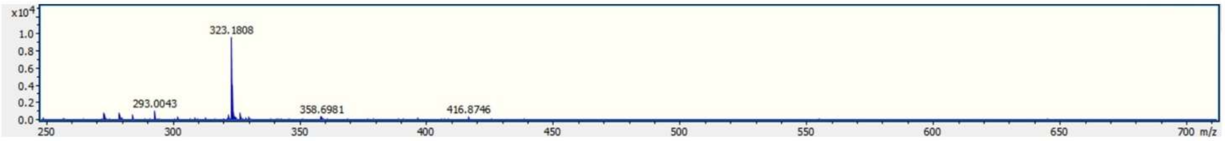

**14e** 50 mM Phosphate, pH 7, 300 mM NaCl, 1 mM CHAPS, 37 °C

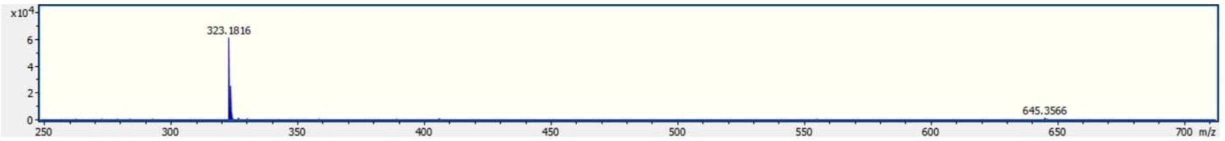

**Figure S14** LC/MS excludes proteolytic degradation of compound **2** by DENV2 protease as an explanation for the weak inhibitory activity in phosphate-based buffers. To understand if the weak activity of compound **2** in phosphate buffer, whether at room temperature or 37 °C, is due to increased degradation by the protease relative to Tris buffer, an experiment was conducted by LC/MS to assess this. Conditions used: DENV2pro 3 $\mu$M, compound **2** 100 $\mu$M, incubation for 24 hr. The non-tagged linked construct was used. Mass of intact compound **2** is $[M+H]^+$ 645 Da and

$[M+2H]^{2+}$ 323 Da. The degradation products with the masses $[M+H]^+$ 407 Da and 257 Da were detected, corresponding to Bz-Arg-Lys-OH and (4-Benzyloxy)Phg-$NH_2$, respectively. For **a** and **b**, incubation of compound **2** (100 $\mu$M) with the protease (3 $\mu$M) for 24 hr at room temperature leads to complete degradation of the compound, where the mass of the intact compound cannot be detected. For **c**, **d** and **e**, the mass of the intact compound is detected at different intensities of 42.1%, 14.9%, and 95.4%, respectively. This shows that the difference in inhibitory activity is not related to the degradation of the compound **2**, but to possibly a different interaction mode with the protein. With the buffers for **a** and **b**, it is expected that the phenylglycine ether extends to the prime side, thus the degradation. However, with the buffers for **c**, **d** and **e**, an alternate binding mode may possibly take place.[1,2] Drazic *et al.* support alternate binding modes for a structure comparable to compound **2** by mass analysis,[3] where in one binding mode the compound is prone to degradation between the phenylglycine ether and lysine but not in the other binding mode.

**Supplementary figures for the non-proteolytic binding assay**

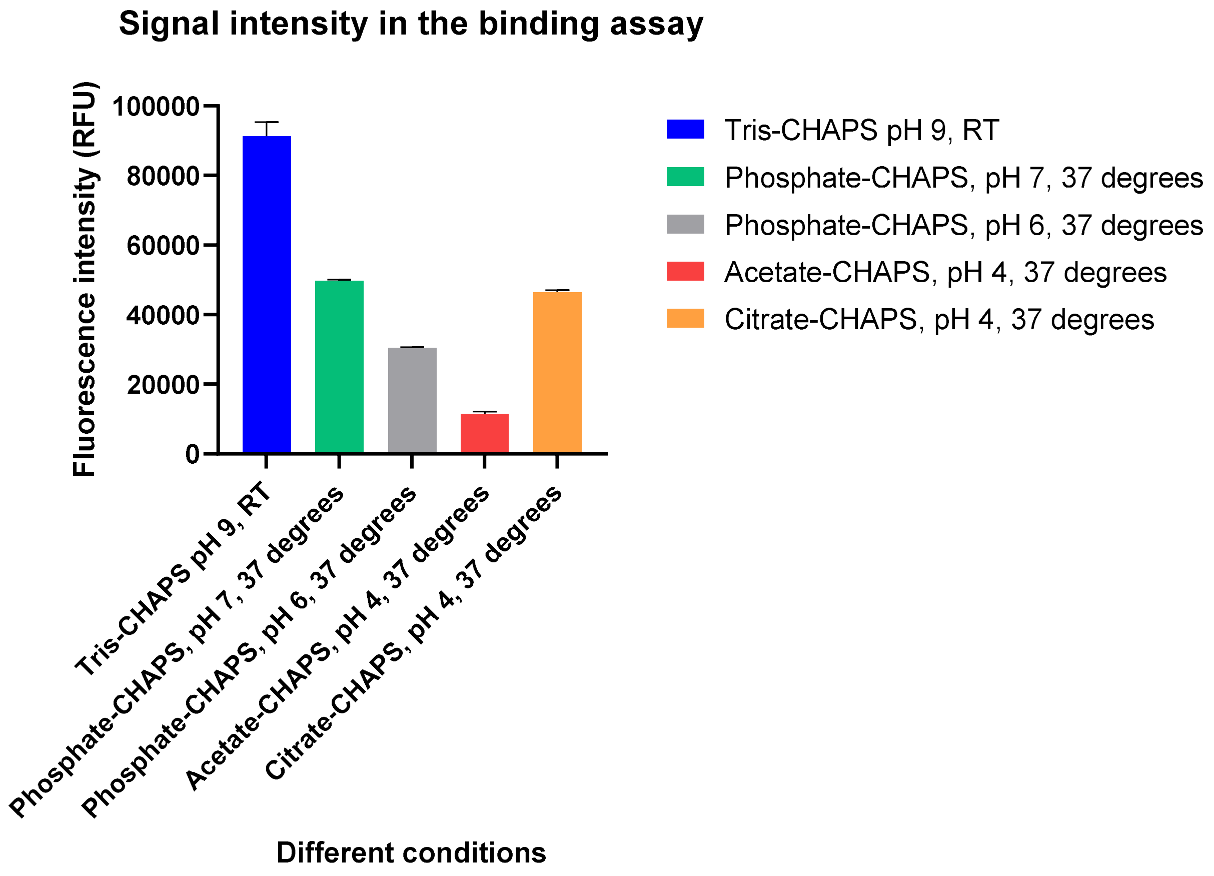

**Figure S15** Signal intensity in the binding assay for 200 nM DENV2 protease. The signal of DENV2pro in the binding assay is shown for different measurement conditions under the same settings for signal amplification, with excitation at 295 nm and emission at 350 nm. For each condition, the incubation was performed for 1 hr. Background signals were subtracted from the recorded fluorescence intensity.

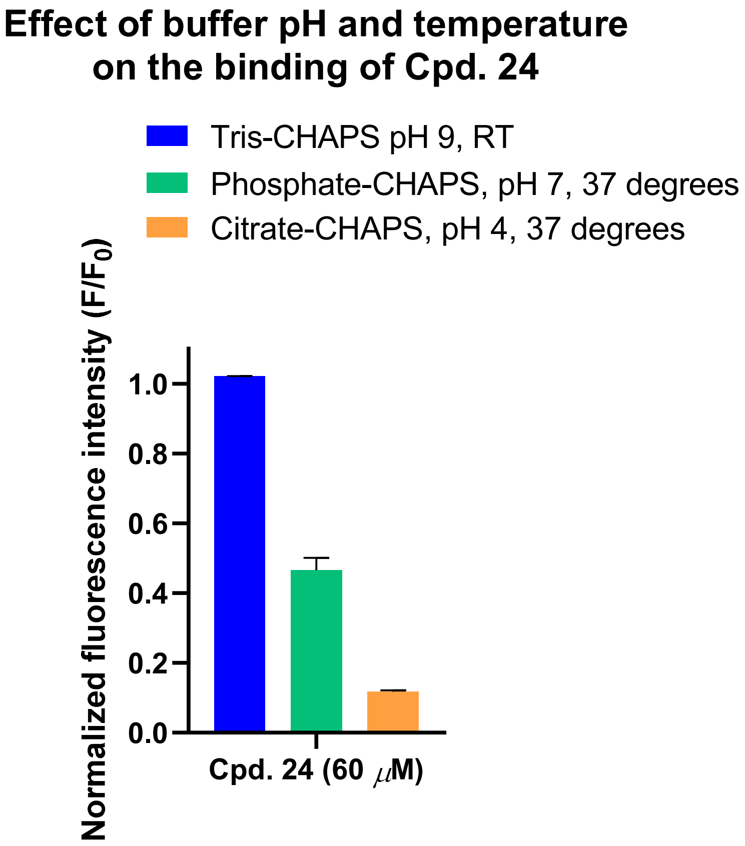

**Figure S16** Effect of buffer pH and temperature on the binding profile of compound **24**. The binding of **24** to DENVpro increases with the decrease in the pH and the increase in temperature. Incubation is performed for 1 hr. Results are corrected for the background and for the inner-filter effects based on the experiments with NATA instead of the protease.

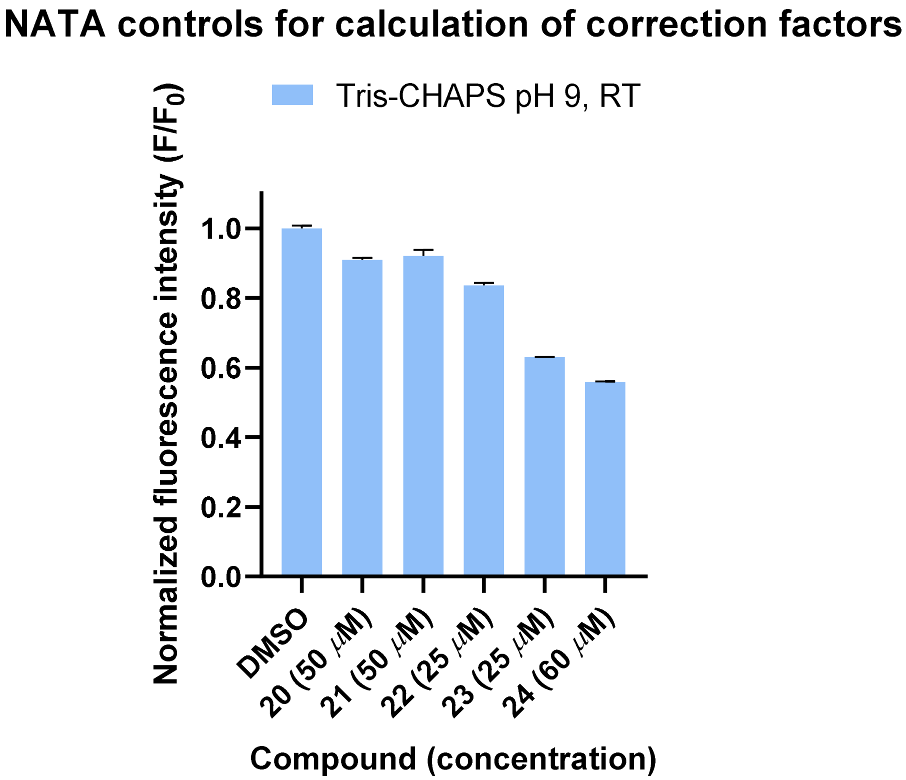

**Figure S17** Representative results of experiments with NATA. The influence of the compounds on NATA (1 $\mu$M) fluorescence instead of DENV2pro show no or limited optical interference by inner filter effects in Tris-CHAPS, pH 9, RT. The observed effects on NATA are used to obtain correction factors for the binding data to confirm specific binding to DENV2pro in the binding assay in Figure 3D.

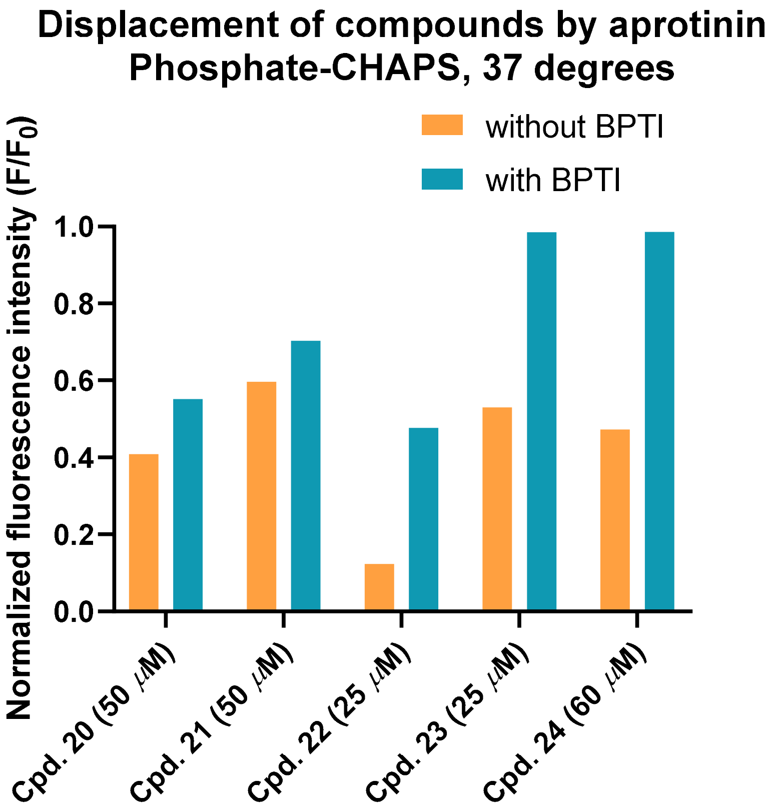

**Figure S18** Displacement of compounds **20-24** by aprotinin (10 $\mu$M) using phosphate-CHAPS buffer, 37 °C and the non-tagged DENV2 protease construct in the binding assay.

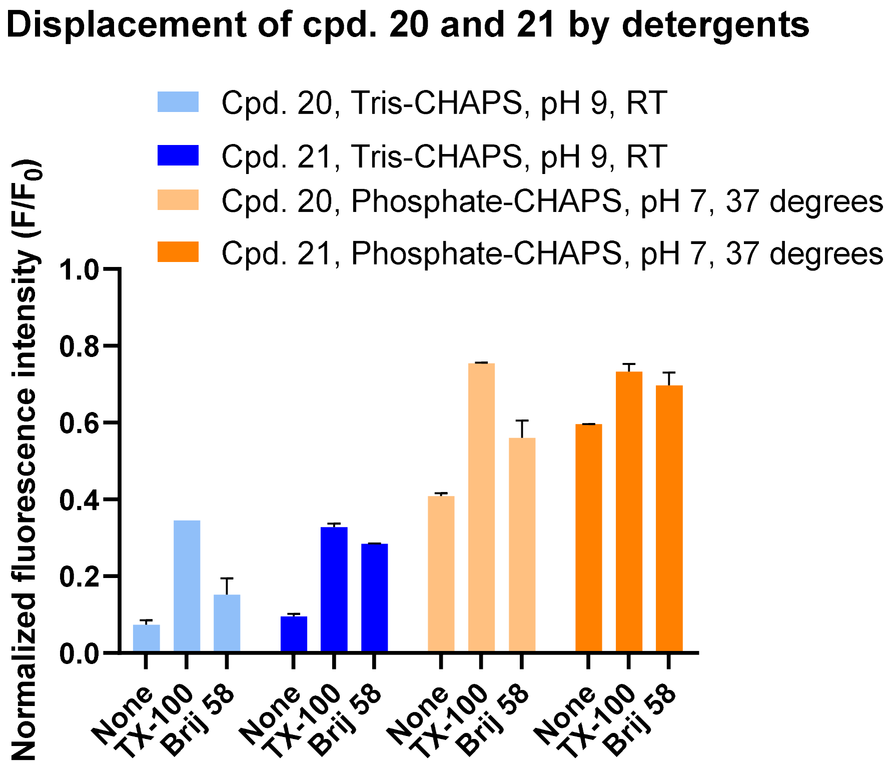

**Figure S19** Displacement of compounds **20** and **21** (50 $\mu$M) in the binding assay by 0.01% (v/v) Triton X-100 or 0.0009% (w/v) Brij 58 using Tris-CHAPS buffer, pH 9, RT, $His_6$-tagged DENV2pro construct or phosphate-CHAPS buffer, pH 7, 37 °C, non-tagged DENV2pro construct.
